# Intrinsic rewards guide visual resource allocation via reinforcement learning

**DOI:** 10.1101/2025.04.25.650663

**Authors:** Ivan Tomić, Rodrigo Raimundo, Paul M. Bays

## Abstract

Humans and other animals prioritise visual processing of stimuli that signal rewards. While prior research has focused on tangible incentives (e.g., money or food), the effects of intrinsic incentives – such as perceived competence – are less well understood. Across a series of visual estimation experiments, we manipulated observers’ subjective sense of confidence in their judgements using either deceptive trial-by-trial feedback or real discrepancies in stimulus reliability. We found that observers prioritised encoding of stimuli associated with lower uncertainty or error, benefiting performance for stimuli already estimated accurately, while further impairing performance for those estimated poorly. These reward-driven biases, while potentially adaptive, impaired overall accuracy in the present tasks by causing resource allocation to deviate from the error-minimizing strategy. To account for these findings, we supplemented a normalization model of neural resource allocation with a simple reinforcement learning rule. Intrinsic and external rewards cumulatively shaped the values assigned to different stimuli by the model, and the resulting discrepancies biased resource allocation and thereby estimation error, quantitatively matching the data. These findings reveal how intrinsic reward signals can shape resource allocation in ways that are both adaptive and counterproductive, offering a computational basis for the motivational biases underlying cognitive performance.

## Introduction

To support adaptive behaviour and ensure survival, the brain has evolved to prioritise environmental cues that signal potential rewards [1, 2]. Selectively attending to reward-predicting stimuli facilitates efficient navigation of complex environments, helping organisms move towards more rewarding states [3, 4]. This selection process is crucial given the brain’s limited processing capacity, as it enhances internal representations of valuable stimuli and facilitates the formation of stimulus-reward associations [5]. Whereas the bias towards processing stimuli associated with tangible rewards is well established, the influence of intrinsic rewards – positive motivational states associated with feelings of satisfaction and competence [6] – on sensory processing remains less understood.

Experiments using points-based and monetary incentives have found that associating stimuli with a higher probability, or greater magnitude, of external reward facilitates voluntary, or top-down, attention [7–9]. Additionally, in visual search tasks, which primarily engage bottom-up processes, search times are faster for pop-out targets associated with higher rewards than stimuli predicting less or no reward [10]. Notably, the prioritisation of reward-associated stimuli persists in subsequent tasks even when reward contingencies are removed, and previously rewarded features cease to be salient or task-relevant [11–13]. Consistent with this, studies have shown that eye movements are biased towards objects and spatial locations previously associated with rewards [14–16]. This continued prioritisation of previously rewarded stimuli, even when it no longer aligns with immediate task goals, suggests that reward learning creates a lasting effect that can involuntarily bias attention towards these stimuli [17, 18].

The influence of external rewards on behaviour extends to visual working memory (VWM) [19], which is known for its ability to flexibly store and maintain features of multiple objects within a limited capacity [20–28]. The precision of representations increases as a function of the associated reward, indicating that VWM allocation also tracks reward values when multiple objects provide different rewards ([29, 30]; see [31] for a review). Objects that were previously associated with reward are also better remembered, even when they are currently task-irrelevant [32]. Crucially, however, total VWM capacity does not show flexibility with reward [33, 34], which is further evidenced by findings that improved performance for high-reward items is accompanied by a corresponding decline in performance for low-reward items [35]. These results demonstrate that stimuli can be strategically prioritised for encoding in VWM through selective attention, leading to flexible allocation of limited capacity between items based on their assigned subjective values [36, 37].

Many everyday behaviours lack immediate external feedback, making internally generated signals – i.e., intrinsic rewards – potentially equally relevant. One hypothesised source of such internal guidance is the subjective sense of accuracy or confidence [38, 39]. Although the neural bases and computational processes underlying intrinsic rewards [6] and metacognition [40] are diverse, recent work suggests that confidence-related signals constitute an important intrinsic driver of learning. Specifically, reinforcement-like learning can emerge solely from an observer’s internal estimate of confidence, used as a proxy for external feedback [41, 42]. The dopaminergic system – and the striatum in particular – which has long been implicated in encoding the motivational significance of actions [43], has also been shown to encode intrinsically generated motivational signals [44], including confidence [42, 45, 46]. Consistent with this, neural signatures typically associated with external feedback are also elicited when individuals experience an internal sense of being correct or incorrect [47–50]. More broadly, the idea that intrinsic signals can support reinforcement learning has a long history in the computational literature [51–53]. The present work identifies confidence-related signals within this framework as a potential driver of attentional allocation.

In the present study, we combined psychophysical measurement and computational modelling to investigate how different intrinsic and external factors affect the competition between visual stimuli for processing resources. We used a modified analogue report task [54, 55] in which observers were instructed to reproduce the direction of one of a pair of motion stimuli that differed in their associated history of reward. Across a series of experiments, we found performance was consistently better for stimuli previously associated with larger external reward, but also those associated with lower uncertainty or with improved performance feedback. To provide a mechanistic explanation of the observed behaviour, we developed a computational model that relates accumulation of past rewards, both intrinsic and external, to allocation of neural resources between stimuli, which in turn influences estimation performance.

## Results

### Differential rewards bias resource allocation

Building on existing evidence that external rewards can bias information processing, we began our investigation by quantifying their effects on representational fidelity in a motion reproduction task. In Experiment 1, observers viewed two coloured motion stimuli, and after a brief delay and the presentation of a colour cue, they were asked to reproduce the motion direction of the cued stimulus (Fig. 1A & B). Critically, in this experiment, we associated the colours of the stimuli with different external rewards by awarding accurate recall (< 50° absolute error) with 15 points when items of one colour were tested versus 5 points for the other colour. Accumulated points were converted into a bonus payment to the observer. At the end of the experiment, all observers correctly identified which stimulus had provided the larger rewards. To determine whether the difference in external rewards influenced reproduction precision, we compared the mean absolute deviation (MAD) of responses between stimuli of different colours. We found strong evidence that response errors were smaller for items of the colour associated with the larger reward (BF_10_ = 18.7, median of the posterior over effect size *δ* = 0.575, 95% credible interval = [0.195, 0.966]) (Fig. 1C & D).

**Figure 1:**
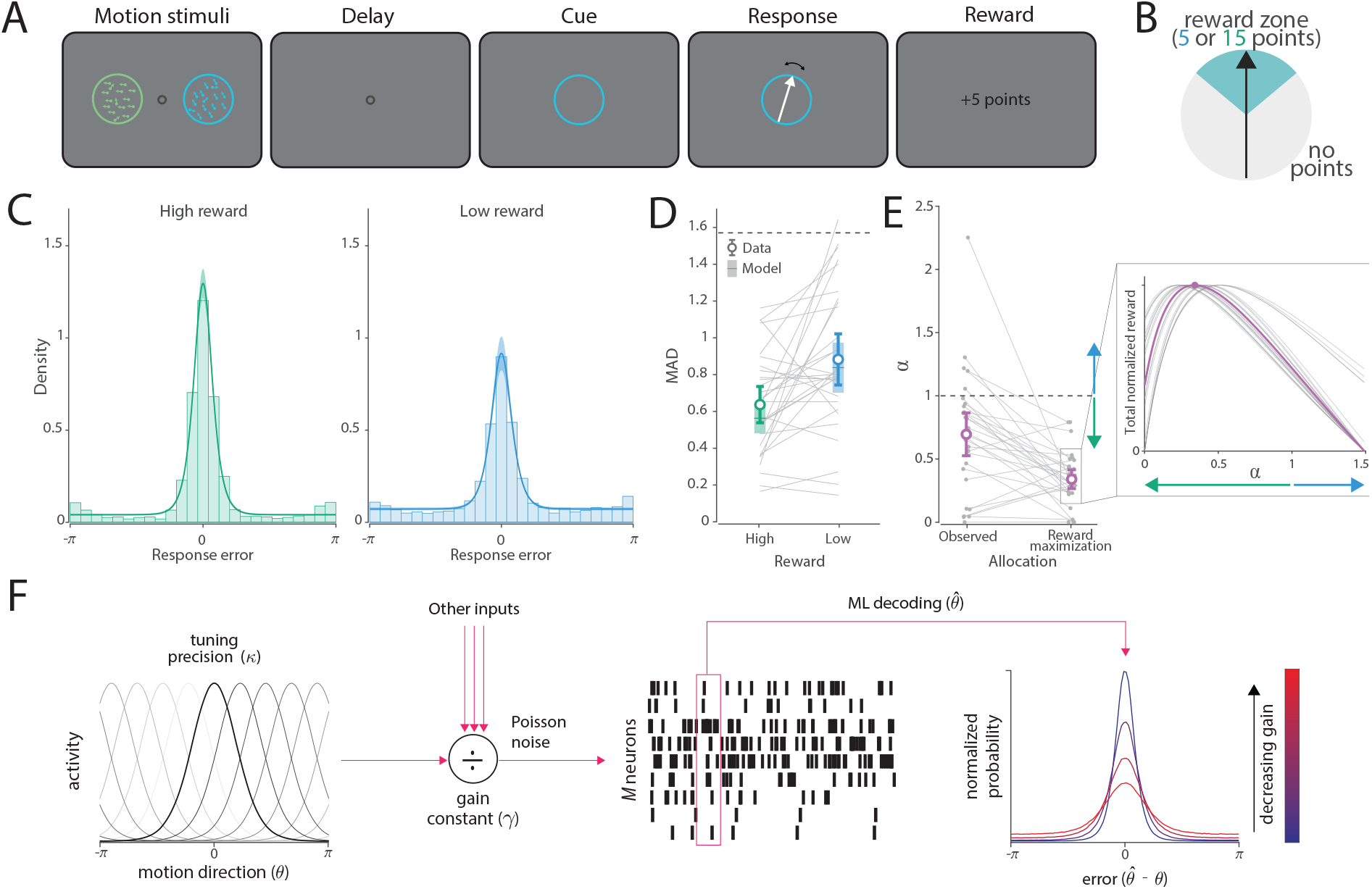
External reward manipulation in Experiment 1. A) Schematic of the task. B) Illustration of the experimental manipulation. Responses within 50 degrees of the target motion direction were rewarded with either 15 or 5 points, depending on the colour of the cued object. C) Distribution of response errors and corresponding fits of the Neural resource model. Histograms represent the data, while coloured curves and shaded areas depict model predictions (M ± SE) D) Corresponding mean absolute deviation (MAD) from experimental data (circles with error bars) and from the Neural resource model (lines with error patches). Error bars and patches indicate the mean and 95% CI. Dashed line indicates chance level performance. E) Observed (i.e., freely estimated) resource allocation compared to the optimal allocation aimed at maximizing the total points in the task. For visualisation purposes, allocation towards the low-reward item is shown. Dashed line indicates equal allocation. Allocation smaller than 1 indicates that more resource was allocated towards the high-reward item (1:0.695 for high- vs low-reward item). Error bars indicate the mean and 95% CI. The inset shows individual reward functions relating resource allocation to expected point totals, with each curve’s peak indicating the allocation that maximizes expected reward. For ease of visualization, only a subset of observers is shown, and all curves are normalized to the same total reward. F) Normalization model of neural resource allocation [22]. Stimulus features are encoded in the activity of a population of idealized direction-selective neurons. All neurons are assumed to have identical bell-shaped (von Mises) tuning curves with concentration *κ*, translated across the feature space to peak at each neuron’s individual preferred feature value, providing dense uniform coverage of the feature space. Each stimulus is encoded by an independent subpopulation of M neurons, and parameter *α* represents a multiplicative gain factor related to attention allocated to each stimulus – and therefore to the activity of each subpopulation (see Methods). Divisive normalization operates across the entire population, scaling summed activity to a fixed level determined by a gain constant. Neurons generate spikes according to a Poisson process, with mean firing rate determined by the normalized input of each neuron. The joint response of the population to a particular stimulus can be visualized as a hill of activity, centred on the true stimulus feature. The subsequent behavioural response was modelled as maximum likelihood decoding of the spiking activity. Examples of error distributions predicted by the Neural resource model (shown right). As the resources allocated to a stimulus (gain) decreases, the distribution of errors becomes increasingly wide and heavy-tailed [56]. when one colour was cued and minified the error at feedback for the other colour (Fig. 2A & B). A post-experimental questionnaire revealed that 84% of observers judged stimuli of the colour associated with error-magnified feedback as more difficult to remember, indicating that we successfully associated stimulus identity (i.e., colour) with perceived difficulty.

#### Neural resource allocation

The results of Experiment 1 indicate that observers prioritised encoding the stimulus associated with the larger reward. Importantly, recall for the low-reward item remained reliably better than chance, suggesting that prioritisation was graded rather than all-or-none. To quantify the share of resources allocated to each item, we applied a normalization-based population coding model [22, 56] to the data from Experiment 1 (Fig. 1F). In this model, neural firing rate takes the role of a limited resource, which, in the simplest scenario, would be equally distributed between stimuli. Here, we extended this model by freely fitting a gain modulation parameter, *α*, which increased the activity encoding one of the two stimuli while keeping the total gain (i.e., mean activity) of the population constant (see the Neural resource model section for more detail). Consistent with the observed difference in response error, we found strong evidence for unequal resource allocation favouring items of the highly rewarded colour (low/high ratio 0.695; difference from equal allocation, BF_10_ = 23.8, *δ* = 0.594, 95% CI = [0.211, 0.987]) (Fig. 1E; (Maximum likelihood (ML) estimates for the other parameters are shown in Supplementary Information).

We next investigated whether observers distributed resources in a way that would maximize the total number of collected points, which we considered an optimal allocation strategy for this task. To test this, we calculated the expected number of points awarded for a range of different allocation weights (see Optimal resource allocation for more detail). The values of *α* that maximized the reward are shown in Figure 1E (optimal allocation). Comparing the observed and optimal weights revealed strong evidence for a difference between the two (BF_10_ = 92.9, *δ* = 0.694, 95% CI = [0.297, 1.102]), with observers distributing resources more equally than would be required to maximize the total number of points (low/high ratio 0.34). Consequently, observers earned fewer points per trial than would be expected under the reward-maximizing allocation (observed = 7.69, optimal = 7.92; BF_10_ = 60.2). Finally, we found moderate evidence against a correlation of observed and optimal allocations (*r* = 0.21, BF_10_ = 0.26).

A reward-maximization strategy would come at the cost of higher error for the low-reward item. This could suggest that, in addition to maximizing external rewards, observers may also aim to achieve a certain level of accuracy on the task across all items, potentially because they find accuracy intrinsically rewarding (see also, [25]).

### Perceived accuracy biases resource allocation

Having confirmed that external rewards modulated allocation in the motion reproduction task, we next investigated effects of perceived accuracy, a possible form of intrinsic reward, on the same task. In Experiment 2a we presented manipulated feedback at the end of each trial to influence observers’ perception of their reproduction accuracy. Studies using false-feedback manipulations have shown that such manipulations can modulate subjective confidence [57], providing a principled basis for our novel trial-by-trial feedback manipulation. Observers were again presented with two coloured stimuli, and reproduced one indicated by a colour cue. We magnified the error presented at feedback

To assess the effects of perceived difficulty on response precision, we compared MAD between the response and the true target direction (rather than the one shown as feedback) for stimuli of the two colours (Fig. 2C & D). We found responses to be more precise for the stimulus with reduced feedback error, i.e., the one perceived as easier to remember (BF_10_ = 29.4, *δ* = 0.672, 95% CI = [0.24, 1.12]). This finding indicates that the perception of better performance for stimuli of one colour, induced by feedback, led to improved actual performance for those stimuli.

**Figure 2:**
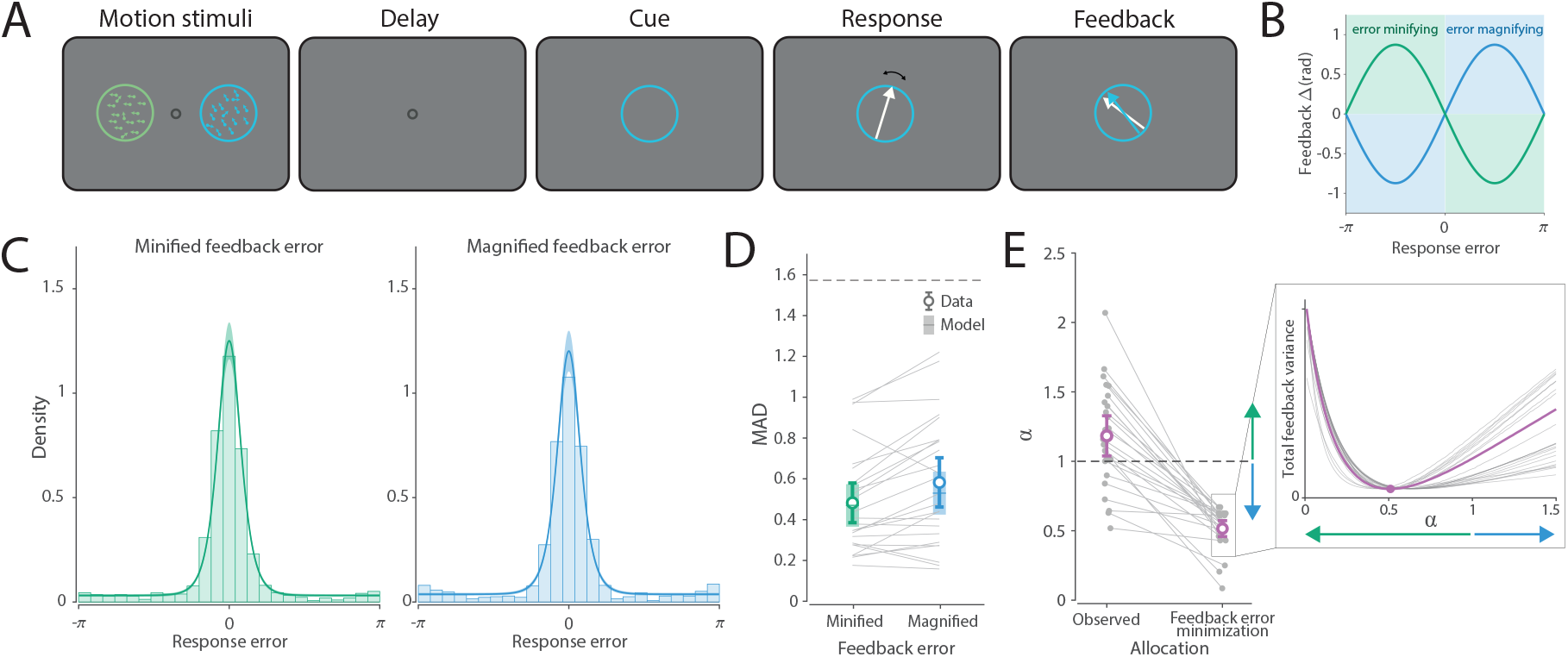
Perceived accuracy manipulation in Experiment 2a. A) Schematic of the task. B) Experimental manipulation illustration. Feedback error was magnified for one stimulus and minified for the other, based on the colour of the cued object. C) Distribution of response errors and corresponding fits of the Neural resource model. Histograms represent the data, while coloured curves and shaded areas depict model predictions (M ± SE). D) Corresponding MAD from experimental data (circles with error bars) and from the Neural resource model (lines with error patches). Error bars and patches indicate the mean and 95% CI. Dashed line indicates chance level performance. E) Observed resource allocation and optimal allocation aiming to minimize overall feedback variance in the task. Dashed line indicates equal allocation. Allocation larger than 1 indicates that more resource was allocated towards the error-minified item (minified vs magnified: 1.18:1). Error bars indicate the mean and 95% CI. The inset illustrates individual variability in feedback error as a function of resource allocation, with each curve’s trough indicating the allocation level that minimizes feedback error. For ease of visualization, only a subset of observers is shown, and all curves are normalized to the same range of feedback variance.

The observed effect could be attributed to either capture of visual attention by the “easier” item (i.e., competition for visual processing resources) or the mnemonic prioritisation of that item (i.e., competition for memory resources). To differentiate between these possibilities, we conducted a follow-up experiment. Experiment 2b replicated the conditions of Experiment 2a but with stimuli presented sequentially to reduce encoding competition between the two objects and minimize the influence of attentional selection on resource allocation. Similar to Experiment 2a, 89% of observers judged the colour associated with magnified feedback errors as more difficult to remember. However, in contrast to Experiment 2a, comparing response errors across the two stimuli (Fig. S1) revealed that the observed data were nine times more likely under the null hypothesis, providing moderate evidence for a lack of difference in response precision between the two colours (BF_10_ = 0.11, *δ* = 0.009, 95% CI = [-0.183, 0.202]). This finding suggests that the effect observed in Experiment 2a is likely due to attentional competition during encoding. When that competition is mitigated, observers do not show preferential encoding based on perceived difficulty.

In addition, we verified whether the results of Experiment 2a could be explained by one object appearing easier to reproduce, the other appearing more difficult, or their combined effect. To this end, we conducted a follow-up, Experiment 2c, which was identical to Experiment 2a except that observers took part in two sessions. In both sessions, feedback for one object was veridical, while feedback for the other object was minified in one session and magnified in the other session. Different colour pairs were used to prevent transfer of learned associations across sessions. Testing the directional hypotheses based on Experiment 2a, we found moderate evidence that the minified item was recalled more accurately than the veridical item (BF_10_ = 7.9, *δ* = 0.557, 95% CI = [0.12, 1.046]; Fig. 3), but no evidence that the magnified item was recalled less accurately than the veridical item (BF_10_ = 0.64, *δ* = 0.253, 95% CI = [0.017, 0.659]). Additionally, we found strong evidence that the minified item was recalled more precisely than the magnified item (BF_10_ = 22.5, *δ* = 0.677, 95% CI = [0.197, 1.046]), replicating the results from Experiment 2a. These results indicate that the error minifying manipulation alone was sufficient to bias resource allocation, whereas the error magnifying manipulation was not.

**Figure 3:**
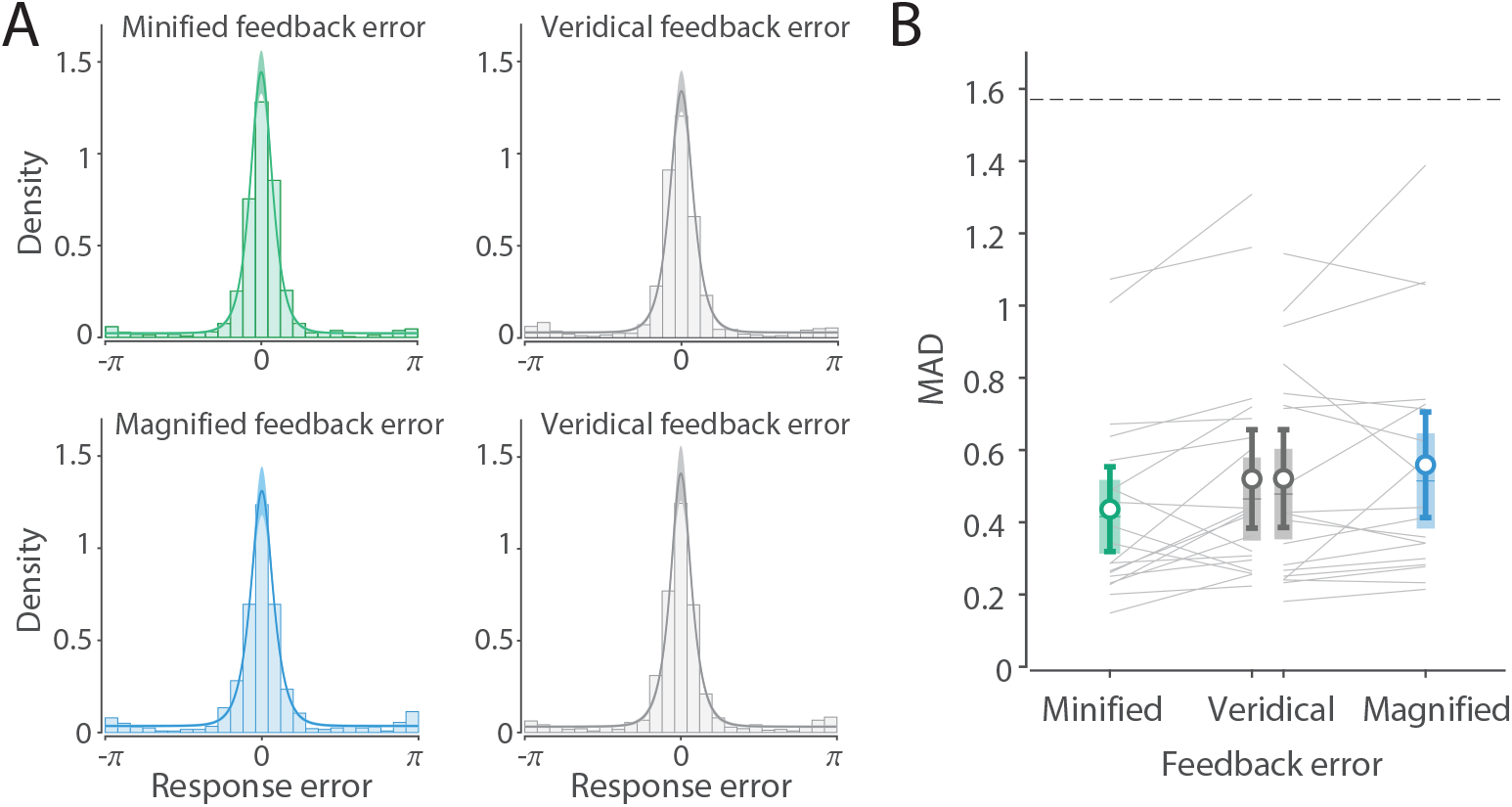
Perceived accuracy manipulation in Experiment 2c. A) Distribution of response errors for the two tasks: in one task, feedback for one object was minified and veridical for the other object (top row); in the other task, feedback for one object was magnified and veridical for the other object (bottom row). Histograms represent the data, while coloured curves and shaded areas depict model predictions (M ± SE). B) Corresponding MAD from experimental data (circles with error bars) and from the Neural resource model (lines with error patches). Error bars and patches indicate the mean and 95% CI. Dashed line indicates chance level performance.

#### Neural resource allocation

The results of Experiment 2a show that observers prioritised encoding of the error-minified stimulus, i.e., the one signalling better performance. Crucially, the error-magnified stimulus was still recalled with above-chance precision, consistent with a graded rather than all-or-none allocation of resources. To quantify resource distribution between the two objects, we again applied the Neural resource model to the data, with results illustrated in Figure 2C & E. We found that, on average, observers allocated 1.18 times more resources towards the error-minified stimulus (difference from equal allocation, BF_10_ = 2.63, *δ* = 0.45, 95% CI = [0.05, 0.86]; 19 out of 25 observers had *α*_*observed*_ > 1; Fig. 2D), consistent with the observed difference in response error between the stimuli of two colours.

Next, we investigated whether the observed allocation matched the predictions of an ideal observer who optimally weights neural activity to minimize overall feedback error in the task. We calculated the expected variance of feedback error across both items for a range of different allocation weights, and Figure 2E shows optimal allocation weights that minimize this variance. The optimal strategy would require shifting twice as many resources towards the error-magnified item (*α*_*optimal*_ = 0.52). Importantly, we found extremely strong evidence that this was inconsistent with the observed allocation, which favoured the error-minified item (BF_10_ = 3.77 × 10^6^, *δ* = 1.72, 95% CI = [1.09, 2.39]). Overall, these results indicate that observers did not adopt an allocation strategy that would minimize their feedback error variability (*α* = 0.52), but instead did the opposite, allocating more neural resources to the item for which we systematically minified the error in feedback. Accordingly, observers’ allocation resulted in higher overall feedback error than predicted by the optimal strategy (MAD_observed_ = 0.535; MAD_optimal_ = 0.476; BF_10_ = 9.92 × 10^3^). We also found moderate evidence against a correlation of observed and optimal allocations (*r* = 0.23, BF_10_ = 0.29).

In Experiment 2b, fitting the same Neural resource model to the data revealed that the observed allocation parameter was numerically close to 1, (*α*_*mean*_ = 1.07; BF_10_ = 0.62, *δ* = 0.18, 95% CI = [-0.01, 0.38]), which aligns with the observed similarity in reproduction precision between the two stimuli. This further supports the conclusion that the effect observed in Experiment 2a depended on attentional competition during encoding. In Experiment 2c, more resource was allocated to the error-minified stimulus compared to the stimulus with veridical feedback (*α*_*mean*_ = 1.21; BF_10_ = 2.1, *δ* = 0.47, 95% CI = [0.022, 0.951]), while resource allocation was found to be comparable between the error-magnified stimulus and the veridical feedback stimulus (*α*_*mean*_ = 0.96; BF_10_ = 0.27, *δ* = 0.11, 95% CI = [-0.305, 0.535]), consistent with the observed pattern of reproduction errors.

### Estimation difficulty biases resource allocation

Following Experiment 2, we aimed to determine whether preferential allocation and encoding would persist when varying objective stimulus difficulty rather than perceived performance. Drawing on previous findings showing a positive correlation in humans between subjective confidence and the motion strength of RDK stimuli [58] (see also [40]), we hypothesised that variations in the objective difficulty of stimuli would modulate internally generated confidence signals, driving the prioritisation of specific stimuli as in Experiment 2. In Experiment 3a, we presented two coloured RDK stimuli with different coherence levels on the majority of trials, to create differences in objective difficulty and associated confidence. We then assessed response precision on the remaining trials, during which both stimuli were presented with equal coherence (i.e., equal difficulty) (Fig. 4A & B).

**Figure 4:**
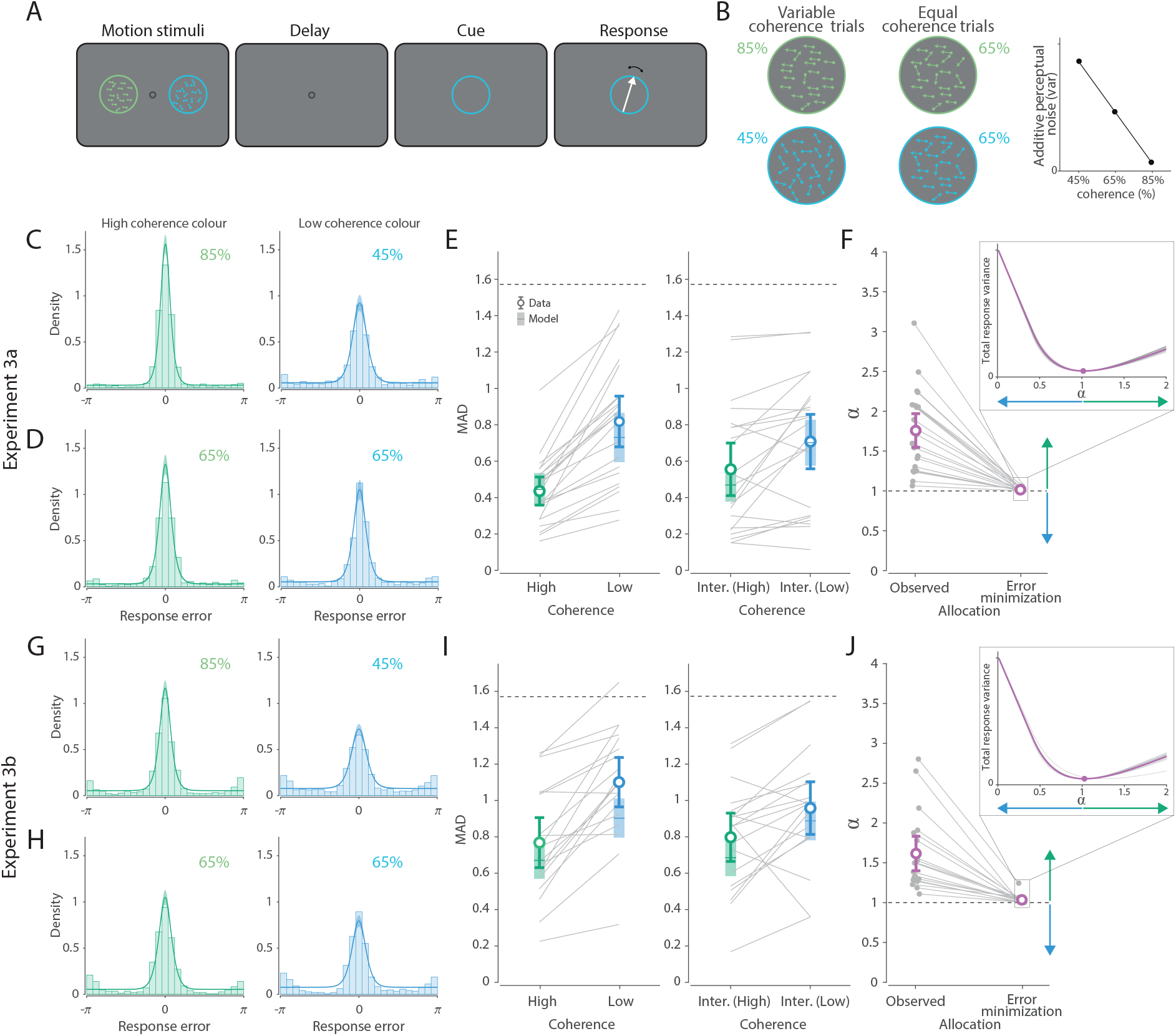
Estimation difficulty manipulation in Experiment 3a and 3b (simultaneous presentation). A) Schematic of the task. B) Experimental manipulation illustration. In most trials, the two colours were associated with different levels of motion estimation difficulty (i.e., variable coherence); in the remaining trials, both objects had the same level of difficulty (i.e., equal coherence). Motion with different coherence levels produces varying degrees of perceptual noise, with higher coherence reducing noise. This perceptual noise was incorporated into the Neural Resource model as an additive component, alongside memory noise. C) & D) Distribution of response errors and corresponding fits of the Neural resource model. Histograms represent the data, while coloured curves and shaded areas depict model predictions (M ± SE). Panel A depicts variable coherence trials, and panel B depicts equal coherence trials. E) Corresponding MAD from experimental data (circles with error bars) and from the Neural resource model (lines with error patches). Error bars and patches indicate the mean and 95% CI. Dashed line indicates chance level performance. F) Observed resource allocation (estimated across all conditions) and optimal allocation (estimated across variable coherence trials) aiming to minimize overall recall variance in the task. Dashed line indicates equal allocation. Allocation larger than 1 indicates that more resource was allocated towards the easier item (high vs low coherence: 1.76:1). Panels G-J are the same as C-F, but for the simultaneous presentation condition of Experiment 3b. J) Allocation larger than 1 indicates that more resource was allocated towards the easier item (high vs low coherence: 1.61:1). The insets illustrate individual variability in response variance as a function of resource allocation, with each curve’s trough indicating the allocation level that minimizes overall response error. Error bars indicate the mean and 95% CI. For ease of visualization, all curves are normalized to the same range of recall variance.

In Experiment 3a, all observers reported that stimuli of the colour associated with low coherence were more difficult to remember, confirming that the coherence manipulation produced a clear difference in perceived difficulty, despite the absence of performance feedback in this experiment. Results of Experiment 3a are shown in Figure 4C-E. As expected, observers were more precise in reproducing the motion direction of the high-coherence stimulus on trials where the stimuli objectively differed in difficulty (BF_10_ = 1.83 × 10^5^, *δ* = 1.6, 95% CI = [0.95, 2.29]). More importantly, on trials where the stimuli had equal coherence, reproduction was also more precise for the stimulus with the colour associated with high coherence (i.e., the “easier” colour) (BF_10_ = 11.4, *δ* = 0.63, 95% CI = [0.18, 1.10]). This finding suggests that observers associated colour with difficulty when the two objects were presented with different levels of coherence, and subsequently allocated more resources to the stimulus they had learned was easier.

This result was replicated in the laboratory setting of Experiment 3b, where observers were required to maintain eye fixation at the centre of the screen during stimulus presentation (Fig. 4G-I). Despite preventing observers from overtly shifting their attention towards one stimulus during encoding, 74% of observers correctly identified one colour as more difficult. As expected, responses were more precise for the high-coherence stimulus on trials where stimuli differed in coherence (BF_10_ = 4414, *δ* = 1.366, 95% CI = [0.718, 2.046]). Additionally, responses were more precise for the colour associated with high coherence on trials where both stimuli were presented with equal coherence (BF_10_ = 4.2, *δ* = 0.56, 95% CI = [0.092, 1.052]). However, the observed difference could again be explained by attentional demands at encoding. When objects were presented sequentially (Fig. S2), response precision was comparable across colours when both stimuli had the same coherence (BF_10_ = 0.51, *δ* = 0.267, 95% CI = [-0.157,0.709]). This was despite observers being able to judge which item was more difficult (84%) and a noticeable precision advantage for the high-coherence stimulus on trials when coherence levels varied between objects (BF_10_ = 9.22 × 10^4^, *δ* = 1.76, 95% CI = [1.015, 2.550]). Compared to other experiments, response distributions in this experiment exhibit more pronounced peaks around the direction opposite to the target (i.e., elevated tail ends), which also produces some exaggeration of MAD compared to the model. The tendency of our sensory system to encode orientation of a motion path (i.e., the line on which movement occurs) partly independently of direction is well-documented [59, 60] and may be especially pronounced when motion stimuli are presented in the periphery rather than at fixation.

#### Neural resource allocation

Consistent with the findings from Experiment 2, Experiment 3 demonstrated that observers, when presented with objects associated with different levels of performance, prioritised the encoding of the stimuli perceived as easier. Also consistent with previous experiments, observers performed above chance for the more difficult item, supporting the interpretation that resource allocation was graded rather than all-or-none. To quantify the distribution of resources across the two items, we again applied our population coding model to the data.

In Experiment 3a, the allocation estimates from the model indicated that observers allocated nearly twice as much resource (1.76:1) to the stimulus colour associated with higher coherence (Fig. 4F), and this allocation deviated from equal allocation (BF_10_ = 2.68 × 10^4^, *δ* = 1.4, 95% CI = [0.792, 2.031]). Interestingly, in this experiment – but not in others – we found evidence that observers with a higher gain parameter on average allocated their resources more equally (*r*_*γ,α*_ = -0.58, BF_10_ = 10.53, 95% CI [-0.78,-0.18]). We next investigated whether the observed allocation was consistent with an optimal allocation strategy aimed at minimizing overall response variance in the task. To this end, we simulated performance on the variable coherence trials using a range of different allocation weights, and found that the optimal strategy for most observers was equal allocation (Fig. 4F). Comparing the observed and optimal weights revealed strong evidence that the observed weights were, on average, larger than the optimal weights (BF_10_ = 9580, *δ* = 1.341, 95% CI = [0.733, 1.977]). This allocation resulted in higher overall response error than predicted by the optimal strategy (MAD_observed_ = 0.627; MAD_optimal_ = 0.569; BF_10_ = 147.8).

Interestingly, our simulation shows almost no variation in estimated optimal allocations (Fig. 4F,J). This shows that allocating resources to the low-coherence item – at least in the current experimental setting – cannot effectively mitigate the effects of perceptual noise introduced by manipulating coherence, without introducing substantial error for the high-coherence item. The only exception to near-equal allocation occurs when the effect of perceptual noise for the low-coherence item is so great as to render estimates effectively random. In this case, resources can be reallocated to improve precision for the high-coherence item without worsening performance for the low-coherence.

These findings were replicated in Experiment 3b. When objects were presented simultaneously, the model estimated that observers allocated resources at a ratio of 1.61:1 in favour of the high-coherence stimulus (Fig. 4J). This allocation was again different from equal allocation (BF_10_ = 830.6, *δ* = 1.166, 95% CI = [0.566, 1.794]), and from optimal, which was again close to equal (mean *α*_*optimal*_ = 1.03; BF_10_ = 789, *δ* = 1.16, 95% CI = [0.561, 1.786]). Finally, fitting a free allocation parameter to the data from the equal coherence condition with sequential presentation (Exp 3b), revealed a ratio of 1.2:1 in favour of the colour associated with high coherence; however, we did not find evidence that this was different from equal allocation (BF_10_ = 0.82, *δ* = 0.343, 95% CI = [-0.09, 0.797]).

### Interim conclusion

In Experiment 1, we observed a clear effect of external rewards on representational fidelity in a motion reproduction task, with observers allocating more cognitive resources to high-reward items. In Experiments 2 and 3, we found a similar effect using novel manipulations, where observers allocated more resources to the stimulus that was perceived as easier, either based on manipulated error feedback (Experiment 2) or internal confidence in estimation (Experiment 3). In both of these experiments, the observed allocation deviated from predictions made by an optimal strategy aimed at minimizing overall feedback or response error. Additionally, we demonstrated that differences in estimation performance were abolished when competition during encoding was removed by presenting stimuli sequentially, suggesting they arise from unequal allocation of attentional resources at the encoding stage.

Building on these results and existing literature [50, 61, 62], we argue that observers in our study found higher accuracy and confidence in their performance intrinsically rewarding, and learned to associate this intrinsic reward with a stimulus feature (i.e., one of the two colours). This association biased resource allocation towards subsequent stimuli with the same feature. Importantly, although reward-driven, this biased allocation was not a strategy that would maximize intrinsic reward on these tasks, because observers had no influence over which stimulus was cued for report on a given trial. Indeed the direction of the biases induced by implicit rewards in Exps 2 & 3 meant that they were counterproductive: increasing overall error variability relative to a strategy of equal allocation. Therefore, instead of evaluating this data from the perspective of optimal performance, we propose a neural model inspired by reinforcement learning to elucidate these findings.

### Reinforcement learning model of resource allocation

To further explore the dynamics of resource allocation, we developed a computational model that integrates principles of neural coding and reinforcement learning. The proposed Reinforcement learning account of resource allocation extends the Neural resource model [22, 56] by incorporating a value-updating mechanism that allows external and intrinsic rewards to influence the future distribution of neural resources (Fig. 5). A key contribution of our model is the concept that rewards – both intrinsic and external – obtained from reproduction of a stimulus become associated with the identifying features of that stimulus, affecting their subjective value and biasing allocation of resources in subsequent encounters. We found that this approach accurately predicted resource allocations estimated by freely fitted allocation weights, indicating that behavioural estimation performance could be successfully inferred from an analysis of accumulated rewards.

**Figure 5:**
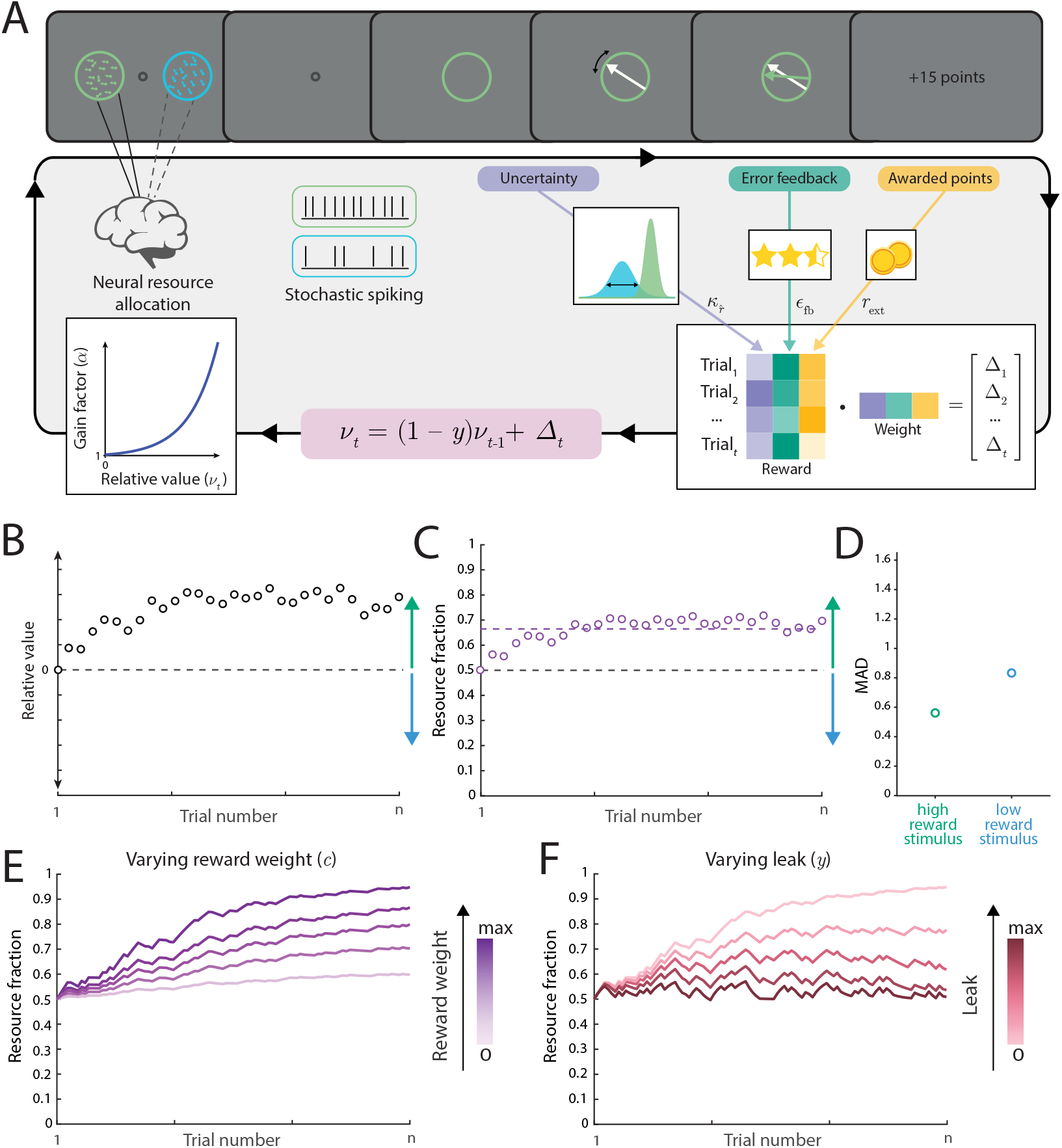
The neural resource allocation account applied to the motion estimation task. A) On each trial, motion directions of the two stimuli are encoded in the spiking activity of populations of neurons, with mean activity determined by the relative allocation of resources to stimuli. Based on the cue colour, one of the populations is decoded to yield an estimated direction with an associated uncertainty that varies from trial to trial. The uncertainty of the estimate, the accuracy feedback (if present) and any points awarded represent different forms of intrinsic and external reward, which are combined as a weighted sum into a composite reward (Δ_t_). This composite reward is then used to update the relative value (*ν*) associated with the stimulus colours. Finally, this relative value is transformed via an exponential mapping into a neural gain factor (*α*), which controls the fraction of resources allocated to each stimulus on the subsequent trial. In this framework, resource allocation is entirely driven by the history of accumulated rewards. B) Throughout the reported experiments, the two colours of stimuli are systematically related to different intrinsic or external rewards, so the relative value assigned to each colour progressively diverges over the sequence of trials. C) Fraction of total resources allocated to the high-reward stimulus over trials, based on relative value shown in B. Note that the fraction of resources allocated to an item is determined by the relative value accumulated up to, but not including, that trial. The dashed line represents the mean allocation across all trials (∼ 65%). The remaining resources (∼ 35%) are allocated to the low-reward stimulus. D) Unequal resource allocation is reflected in differences in the mean absolute error across trials when the high- or low-reward stimulus is cued for report. E) Larger reward weight (unspecified reward *c*), with a constant leak factor, results in a stronger preference for one stimulus over the other in terms of resource allocation. F) Larger leak factor (*y*), with a constant reward weight, leads to a weaker preference.

#### External reward

In Experiment 1, the stimulus colour associated with a high reward (15 points) was expected to accumulate greater value relative to the colour associated with a low reward (5 points). On average, observers earned points on 80% of trials when the high-reward stimulus was probed and 66.5% of trials when the low-reward stimulus was probed, leading to an average accumulation of 602 and 166 points, respectively. To apply the proposed RL model to each observer’s data, we combined individual trial-by-trial external rewards with estimates of internal confidence (Equation 9).

Figure 6A shows the average trajectory of resource allocation across trials (see Fig. S5A for individual trajectories). This trajectory shows an early shift in resource allocation towards the preferred item, followed by a stable plateau. For ease of visualisation, trajectories are presented as directed towards the preferred object, defined as the object receiving a greater average resource allocation across all trials.

**Figure 6:**
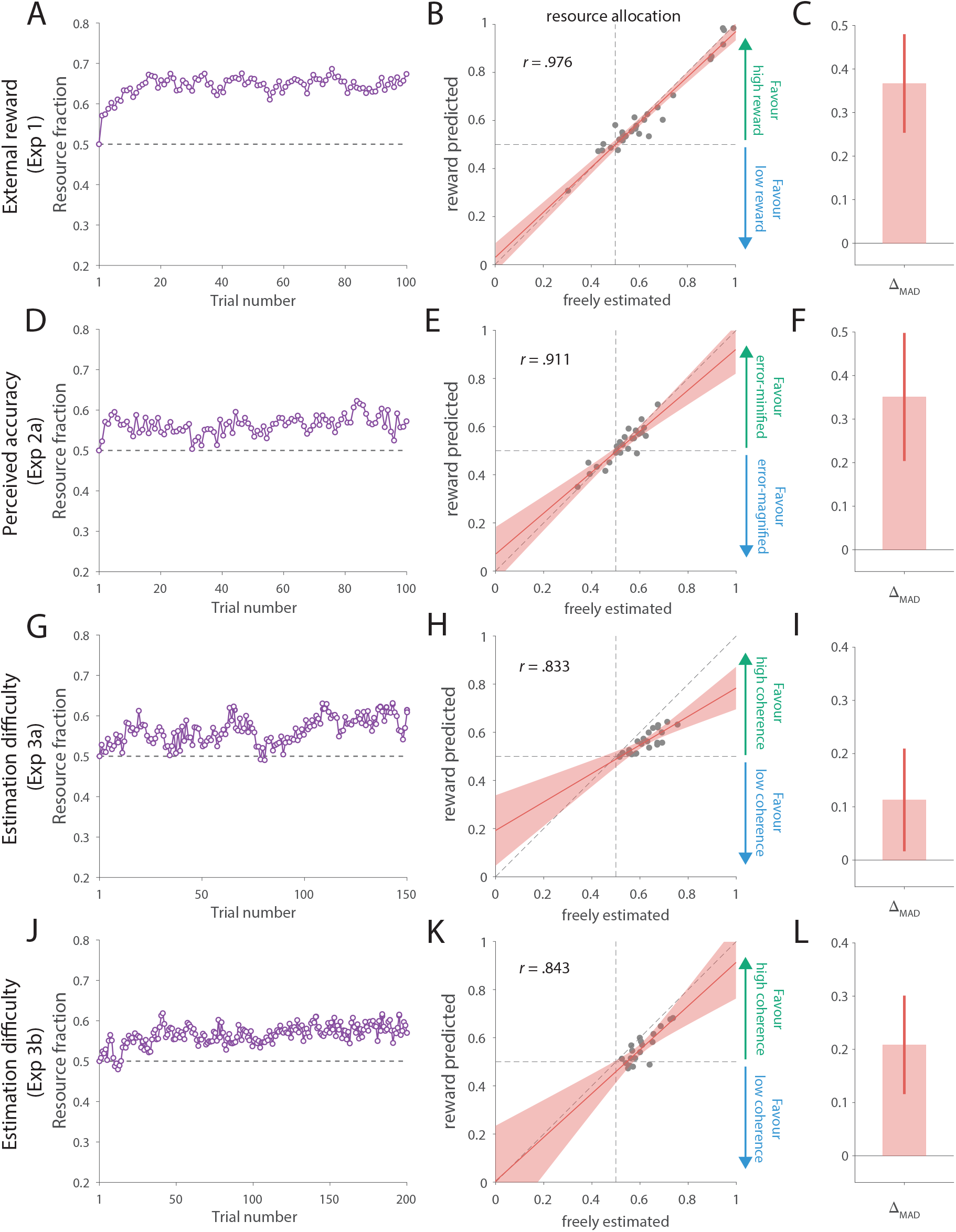
Modelling results. A) Resource allocation across trials inferred by the RL account in the external reward experiment (Experiment 1). Circles represent the mean fraction of resources across observers allocated on each trial towards the overall preferred object. B) Correlation between mean allocations inferred by the RL account and freely estimated allocations. The red line shows predictions of the fitted linear regression model, and the shaded area indicates the 95% CI. C) Difference in MAD between trials on which the probed item had below- and above-median resources allocated to it, as estimated by the RL account. On average, MAD was larger when less resource was allocated to the probed stimulus. Error bars show the mean and 95% CI. D–F) Same as above, but for the perceived accuracy experiment (Experiment 2). G–I) Online estimation difficulty experiment (Experiment 3a). J–L) Lab-based estimation difficulty experiment (Experiment 3b, simultaneous condition).

Crucially, since our RL account is grounded in the same Neural resource model previously employed to fit the psychophysical data and quantify resource allocation (Fig. 1A & C), we can directly compare estimates across the two models. Here we focus on the comparison of estimated resource allocations, while ML estimates and comparisons for the other parameters are shown in Supplementary Information (Fig. S6A). Importantly, the freely estimated resource allocation (observed allocation in Fig. 1C) is based on behavioural errors only, with no information about rewards, and so can serve as a benchmark for evaluating performance of the RL model. As shown in Figure 6B, we observed a strong positive correlation between the freely estimated allocation parameter and the mean allocation derived from the history of accumulated rewards (*r* = 0.976, 95% CI = [0.941, 0.988], BF_10_ = 3.34 × 10^16^). The close alignment of these two distinct methods suggests that the history of accumulated rewards can effectively account for resource allocation in this task.

#### Intrinsic reward: Perceived accuracy

In Experiment 2 we manipulated the response error presented in feedback to influence the perceived difficulty of reproducing stimuli of each colour. This manipulation resulted in participants experiencing systematically larger feedback errors for stimuli of one colour (magnified feedback MAD = 0.837) than the other (minified feedback MAD = 0.233). To model this data within our RL account, we assume that the feedback on each trial provided an intrinsic reward that was associated with the corresponding stimulus colour. This assumption is supported by evidence that observers’ subjective evaluations tend to favour smaller feedback errors over larger ones because they suggest higher accuracy [63].

In the model, rewards derived from feedback were integrated with those derived from internal confidence. Figure 6D illustrates the mean trajectory of resource allocation across trials (Fig. S5B shows individual trajectories for example observers). The model fits again indicate that resources were unequally allocated between stimuli, although the bias is smaller than observed in the experiment with external rewards.

Comparing the estimated allocation derived from the RL account to the freely fitted allocation parameter in the Neural resource model (Fig. 6E), we found a strong positive correlation (*r* = 0.911, 95% CI = [0.777, 0.958], BF_10_ = 3.44 × 10^7^). Consistent with the findings from Experiment 1, the correspondence between these two distinct approaches indicates that the history of accumulated intrinsic rewards provides an explanation for resource allocation in the task with manipulated feedback. ML estimates and comparisons for the other parameters are shown in Supplementary Information (Fig. S6B).

#### Intrinsic reward: Estimation difficulty

Experiment 3 investigated the role of objective difficulty in the representation of motion information. On most trials, two stimuli with different coherence levels (85% and 45%) were presented. We hypothesized that internal confidence in each item’s motion direction, reflecting a metacognitive estimate of accuracy, functions as an intrinsic reward which observers associate with each item’s identity (i.e., colour) [42]. To model the psychophysical data in the simultaneous presentation condition, we estimated internal confidence by exploiting the close coupling between uncertainty and trial-to-trial variability in error within the Neural resource model. Informed by observed response error on each trial, we derived the posterior probability distribution of likelihood precision and used the most probable precision as a basis for intrinsic reward (Eq. 12). While internal confidence was also incorporated in this way when modelling data from the previous two experiments, in Experiment 3 it was the sole source of reward influencing resource allocation.

Figure 6G & J show the mean trajectories from Experiment 3a & 3b, respectively. Again, we visualised the obtained individual trajectories in example participants (Fig. S5C & D). In both experiments, all parameter estimates obtained with the Neural resource model and the RL account strongly covaried (Fig. 6H & K & Fig. S6C & D). Importantly, this was also true for estimates of resource allocation. Across the two experiments, we found very consistent and strong positive correlations between the freely estimated allocation parameter and the mean allocation derived from the history of accumulated rewards (Exp3a: *r* = 0.833, 95% CI = [0.589, 0.923], BF_10_ = 1.29 × 10^4^; Exp3b: *r* = 0.843, 95% CI = [0.575, 0.933], BF_10_ = 3.67 × 10^3^). Consistent with the findings from the first two experiments, this strong correspondence indicates that the history of accumulated intrinsic rewards based on internal confidence effectively accounts for resource allocation in this task.

#### Changes in resource allocation predict response precision

Our finding that freely estimated resource allocation strongly correlates across participants with resource allocation based on the history of rewards supports the conclusion that human resource allocation is guided by a reward-driven value assignment to objects in the visual environment. To further substantiate this claim, we investigated whether variability in resource allocation across trials within individual participants, derived from the RL model, predicts the magnitude of their response errors.

To investigate this, we examined response errors as a function of model-predicted resource allocation, with the hypothesis that trials with lower allocated resources to the probed item would show greater errors. To this end, we performed a median-split analysis for each observer based on the estimated fraction of allocated resources towards the item associated with larger reward (i.e., individual trajectories similar to those shown in Fig. 6, left column). Specifically, we calculated the MAD of response errors for trials with above- and below-median resource allocation, separately for trials where the high- or low-reward item was tested. We hypothesised that MAD would be greater on trials where the RL model indicated that below-average resource was allocated to the probed item, i.e., below-median trials when the high-reward object was probed and above-median trials when the low-reward object was probed. To test this, we computed a composite score for each observer equal to the sum of the signed difference in MAD between below- and above-median trials when the high-reward item was probed and the signed difference in MAD between above- and below-median trials when the low-reward item was probed. In all four experiments, the composite scores indicated that lower predicted resource allocation, based on the history of rewards, corresponded on average to larger MAD of response errors (Fig. 6C, F, I & L) This was confirmed with one-sided t-tests against zero which provided moderate to extreme evidence for a difference in the hypothesised direction: Experiment 1 (BF_10_ = 5.55 × 10^4^, *δ* = 1.096, 95% CI = [0.635, 1.570]); Experiment 2 (BF_10_ = 564, *δ* = 0.866, 95% CI = [0.402, 1.345]); Experiment 3a (BF_10_ = 3.74, *δ* = 0.441, 95% CI = [0.073, 0.874]); Experiment 3b (BF_10_ = 192.4, *δ* = 0.919, 95% CI = [0.377, 1.487]).

## Discussion

In the present study, we investigated how human observers represent stimuli associated with varying levels of external and intrinsic reward. Across three psychophysical experiments, we paired object identities with different rewards and found observers developed higher estimation accuracy for the items associated with larger rewards. In two additional experiments, we demonstrated that this effect was driven by competition for attentional, rather than mnemonic, resources. To provide a mechanistic explanation of this behaviour, we developed a neural model incorporating a reinforcement learning rule that directs resource allocation towards more rewarding stimuli. Our key finding – based on our experimental data and computational modelling – is that a resource allocation mechanism based solely on the history of accumulated rewards is sufficient to explain differences in estimation performance based on intrinsic as well as external rewards.

In the first experiment, we investigated the effects of external rewards on representational fidelity in a motion reproduction task. Both the psychophysical results and computational modelling provided compelling evidence that observers allocated more processing resources to objects associated with a higher reward, resulting in more precise reproduction of high-reward stimuli compared to low-reward ones. This finding aligns with a broad body of research demonstrating that external rewards, such as points or money, influence various aspects of information processing, including the allocation of attentional resources [17, 18] and working memory [31], while also facilitating motor responses, such as hand movements and saccades, towards rewarding stimuli [63–65].

Although intrinsic motivation has long been recognised as a component of reinforcement learning [51–53], its influence on representational fidelity in perceptual and memory tasks has received comparatively little empirical attention. Building on the premise that accuracy itself is rewarding [50, 61, 62], we conducted two experiments that manipulated perceived accuracy (via feedback) and objective estimation difficulty (via signal strength) in a motion reproduction task. We found converging evidence at both the behavioural and computational levels indicating that observers allocate more neural resources towards objects associated with better estimation performance – this in turn improves the internal precision of the corresponding object when encountered on the next trial, which further reinforces the association between object identity and reward. This was the case whether performance differences were induced by experimentally manipulated feedback (Experiment 2) or by objective differences in stimulus discriminability (Experiment 3).

We argue that observers derived intrinsic reward from confidence in their responses and feedback on their accuracy. In our tasks, the association of these rewards with the distinguishing feature of the presented objects (i.e., colour) leads to a bias in resource allocation, favouring subsequent stimuli that share the same feature. This proposal aligns with the notion that perceptual features linked to rewards are prioritised in sensory processing due to their incentive salience (e.g., [66, 67]). Moreover, neural evidence supports this notion by demonstrating that sensory representations are modulated by the history of rewards, underscoring the impact of reward associations on perceptual processing [68]. To make our proposal concrete, we developed a mechanistic model grounded in the principles of population coding and reinforcement learning. Specifically, our reinforcement learning account operates by analyzing accumulated rewards and allocating proportionally more resources to objects previously associated with higher rewards. We found this model closely replicated resource allocation estimates obtained from freely fitted parameters, suggesting that the history of accumulated intrinsic and external rewards is sufficient to account for the observed patterns of resource allocation.

A key novel finding from the proposed model is that both internally generated and externally manipulated (via feedback) estimates of accuracy, when associated with an object’s distinguishing feature, can bias the subsequent processing of objects that share that feature. While the role of external feedback in reinforcing behaviours more generally has been widely acknowledged (e.g., [69, 70]), recent research demonstrates that internal confidence can similarly reinforce behaviour even in the absence of explicit feedback. For instance, improvements in sensitivity in a perceptual learning task have been observed without external feedback [71]. Guggenmos et al. [42] proposed that such learning is driven by confidence prediction errors – discrepancies between an individual’s current confidence and their expected confidence level. Notably, the neural substrate for these prediction errors has been identified in the striatum (see also [72]), a brain region traditionally linked to reward processing. Our findings contribute to a growing body of literature that highlights the importance of metacognition [40] and self-reinforcement [38] as critical processes in the pursuit of rewards.

In the present experiments, we did not measure confidence directly; instead, we relied on manipulations that have been shown to selectively alter subjective confidence [57, 58, 73, 74], together with computational modelling that tracks internal uncertainty without requiring explicit metacognitive reports [56, 75]. These aspects of our design allowed us to probe confidence-related processes without imposing additional metacognitive demands. In support of this interpretation, false-feedback manipulations have been shown to reliably modulate subjective confidence while leaving accuracy unchanged [57, 73, 74]. Given these findings, the effects we observe under our trial-by-trial feedback manipulation are unlikely to be explained solely by expected performance derived from observing feedback, and instead point to changes in internal confidence. Nevertheless, recent work has underscored the importance of more clearly dissociating expected accuracy or success from confidence when examining the mechanisms that guide (meta)cognitive performance. Monetary-incentive paradigms achieve this by manipulating reward valence (i.e., gains vs losses; [76, 77]. Future work should build on these observations by explicitly orthogonalizing accuracy and confidence signals across other types of task scenarios used here. Additionally, although high accuracy in detecting which item provided greater reward prevents us from examining individual differences in allocation as a function of observers’ beliefs about reward, future studies could more directly investigate how expectations about differential rewards, feedback, and difficulty influence resource allocation. Finally, due to their relatively low frequency, we did not explicitly model the 180-degree errors commonly observed in motion perception, whose origin has been discussed in previous work [59, 60]. An important open question is whether these errors in the present setting are indicative of attentional resource allocation, and specifically whether they increase when attentional resources are shifted towards another item.

Using a range of tasks similar to the one used in this study, previous research has demonstrated that humans possess knowledge about the uncertainty with which individual items are reported (e.g., [58, 78, 79]). Population coding models [80, 81] have been particularly effective in capturing subjective confidence [75], as well as proxies such as response latency [82]. Within the population coding framework, an ideal observer of spiking activity would derive their confidence estimate – whether internal or explicitly reported – based on the precision of the posterior distribution, which represents the probability of the stimulus value given the observed neural activity. In the Neural resource model [22, 56], the precision of the posterior (or likelihood, assuming a uniform prior over stimulus space) varies from trial to trial, as a result of stochastic variation in the number of spikes available for decoding. We calculated the most probable estimate of posterior precision on each trial to serve as an indicator of internal confidence. On this basis, the model successfully recreated freely estimated resource allocations based on our data.

We estimated the optimal allocation strategy that would maximise external reward and expected accuracy (or equivalently, minimise feedback and response error in Experiments 2 and 3, respectively) and simulated performance under this policy. Across all experiments, the observed allocations systematically deviated from this optimal pattern, and observers’ achieved outcomes – number of points, feedback error, and recall error – were consistently worse than those predicted under optimal allocation. These observations indicate that participants did not behave in a way that would maximise expected accuracy. For their behaviour to nevertheless be strategic, the resulting policy would have to coincidentally resemble that expected under an intrinsic, implicit reinforcement mechanism, while simultaneously being suboptimal – and even detrimental – with respect to accruing rewards.

In a similar vein, a recent study [83] provided theoretical and empirical evidence suggesting that sensory processing is optimised to maximize fitness (i.e., rewards), rather than to ensure perceptual accuracy. Supporting this idea, neurophysiological studies have demonstrated that early sensory systems encode both sensory information about a stimulus and non-sensory information regarding the behavioural relevance of stimuli [3, 84]. Embedding stimulus-reward contingencies within the sensory representation of a stimulus facilitates the prioritisation of behaviourally relevant information during encoding. These previous findings may help explain why our observers’ allocation strategies were not optimized for accuracy in the task, however they were also not optimized for maximizing rewards. In the experiment with external rewards we found that observer’s allocated resource more equally between items than would be predicted by a reward-maximizing strategy. The RL model captured the observed allocation strategy based on a weighted combination of points-based external rewards and confidence-based intrinsic rewards – this combination of factors could lead observers to maintain a certain level of performance even for stimuli associated with low external reward. When considered across all experiments, our results point to a reward-driven allocation of resources that, while prioritising reward-related stimuli, is not optimized to obtain rewards in the specific tasks we investigated.

In the broader literature on confidence and intrinsically motivated behaviour, it is well established that information-seeking (i.e., curiosity) tends to peak at intermediate levels of confidence [85]. This reflects a general tendency for people to disengage from tasks they are either unlikely to succeed at or highly confident they can solve. Similar effects have been observed in attentional allocation: for example, infants preferentially attend to events with intermediate levels of information [86, 87]. Delayed reproduction of analogue features, such as motion direction in the present study, is a challenging task even with low-noise high-contrast stimuli. However, we cannot rule out the possibility that in a simpler task where the response associated with one item could be made sufficiently trivial, a similar disengagement of attention and reallocation to the more challenging item might be observed. An important question for future studies will be how intrinsic reward signals, including confidence, influence resource allocation within the broader range of intrinsically motivated behaviour.

Our results also contribute to prominent theories in neuroscience, psychology, and economics [88–91] which consider how humans and other animals link the mental effort required for a task with the value of its outcome (i.e., the reward). Behavioural studies demonstrate that, when faced with tasks offering equal rewards but varying in effort, humans tend to avoid those perceived as more difficult [92, 93]. Based on this, it has been argued that cognitive effort is experienced as carrying disutility, i.e., acting as a discount factor on expected rewards [91, 94]. This hypothesis has been substantiated by the observation that cognitive effort reduces neural responses to rewards following an effortful task [95]. In the present results, perceived (Experiment 2) or objective difficulty in estimation (Experiment 3) similarly appears to have discounted or reduced the subjective value of a stimulus, leading observers to prioritise *easier* – and thus in principle more *rewarding* – items for encoding. However, because observers had no control over which stimuli were selected for test, this allocation strategy did not result in more reward in our tasks and could even be counterproductive. This raises the wider question of whether humans may similarly allocate effort suboptimally, driven by intrinsic reward, in other situations where they have limited control over what information will subsequently become relevant.

## Materials and methods

### Apparatus

In the online experiments, tasks were presented via web browsers on observers’ personal computers and were coded in JavaScript and HTML Canvas. In the laboratory experiment, stimuli were displayed on a 69 cm gamma-corrected LCD monitor with a refresh rate of 60 Hz. Observers were seated in a dark room and viewed the monitor from a distance of 60 cm, with their heads supported by a forehead and chin rest. Eye position was monitored online at 1000 Hz using an infrared eye tracker (SR Research). Stimulus presentation and response registration were controlled by a script written in Psychtoolbox [96, 97] and executed in Matlab (The Mathworks Inc.). Responses were collected using a computer mouse.

### Participants

A total of two hundred fifteen naive observers (118 females, 91 males, 6 preferred not to say; M_*age*_ = 27.7, SD_*age*_ = 5.0) took part in the study after giving written informed consent in accordance with the Declaration of Helsinki. All observers reported normal colour vision and normal or corrected-to-normal visual acuity. For the online experiments, observers were recruited using Prolific (https://www.prolific.co) and were remunerated £6 per hour for their participation. For the laboratory experiments, observers were recruited through the Cambridge Psychology research sign-up system and were remunerated £10 per hour. Observers in Experiment 1 received a bonus payment proportional to the number of collected points. In contrast, observers in Experiments 2 and 3 did not receive any additional bonus payments. Our rationale was to avoid introducing external reward contingencies into experiments that were intended to be compared against Experiment 1.

For the online experiments, we used a Bayesian stopping rule to determine the sample size. The stopping rule guides when enough evidence has been gathered to support a decision, thus optimizing the sample size. In particular, we continued testing observers until we obtained strong evidence, as estimated by the Bayes Factor, in favour of either *H*_0_ (BF_10_ ≤ 0.1, indicating evidence supporting no difference between the two conditions of interest) or *H*_1_ (BF_10_ ≥ 10, indicating evidence supporting a difference between the two conditions). If neither hypothesis was supported, data collection ceased after reaching 100 observers. In Experiment 1, we assessed differences in mean absolute reproduction error in the analogue report task between stimuli associated with high and low reward, which were the conditions of interest for the Bayesian stopping rule. In Experiment 2, we tested for differences in mean absolute reproduction error between error-minified and error-magnified stimuli (Experiments 2a and 2b), and additionally included veridical items in Experiment 2c. In Experiment 3, we compared mean absolute reproduction errors on trials where stimuli were presented in different colours but with equal coherence. For the laboratory experiment (Experiment 3b), we aimed to collect a number of participants similar to that in Experiment 3a. In total, thirty observers participated in Experiment 1. Twenty-five observers participated in Experiment 2a, one hundred participated in Experiment 2b, and nineteen participated in Experiment 2c. Finally, twenty-two and nineteen observers participated in Experiments 3a and 3b, respectively.

### Stimuli

The stimuli in this study were random dot kinematograms (RDK). On each trial, two RDK stimuli, each consisting of 40 dots, were presented within a circular aperture. A percentage of the dots (specified below) moved in a coherent direction, while the remaining dots moved in random but consistent directions within the aperture [98]. When a dot exited the aperture, it was replaced by a new dot at the aperture’s edge, maintaining a constant dot density. In all experiments, one stimulus was always green (RGB colour values; online: 47, 195, 129, lab: 0, 199, 128) and the other was always blue (online: 24, 199, 233, lab: 0, 187, 241). In Experiment 2c and Experiment 3b, the same observers completed two tasks (see below). In these experiments, stimuli were either green and blue or orange (237, 154, 0) and magenta (255, 79, 208), balanced across observers and presentation conditions. Across all tasks, stimuli were presented against a mid-grey background. The motion direction for each stimulus was drawn randomly and independently on every trial.

For the online experiments, all measures in pixels are reported for a 1920 x 1080 resolution and 60 Hz refresh rate. When a different resolution or refresh rate was detected, all measurements of size, positioning and speed were automatically adjusted to maintain consistency in stimuli presentation across different display settings. The stimulus aperture was 105 pixels in diameter, and each dot had a radius of 3 pixels. Two apertures were positioned 220 pixels to the left or right of the screen centre. On each frame, the dots were shifted by 3 pixels in a specific direction. In the laboratory experiment, two apertures (1.4 dva radius) were presented horizontally aligned with the fixation annulus, positioned at 5 dva to the left and right. Each dot was 0.15 dva in diameter and travelled at 4 dva/sec speed.

### Procedure and task

In all experiments, observers completed an analogue report task [55]. Each trial began with the presentation of a central fixation annulus. In the laboratory experiment, gaze direction was monitored using an eye-tracking camera, and observers were required to maintain gaze fixation within a radius of 2° around the central annulus for 500 ms before the trial could proceed. After achieving stable fixation, the fixation annulus changed appearance (i.e., became thinner) to signal that the memory array would be presented in 500 ms. In the online experiment, the appearance of the fixation annulus changed after a fixed interval of 500 ms. The sample array, consisting of two RDK stimuli, was then shown for 750 ms, followed by a 1000 ms delay period. A centrally presented colour cue subsequently indicated which of the previously presented stimuli, distinguished by colour, was the target that the observers should recall and report the direction of.

Once observers were ready to give their response, they could begin moving the cursor with a mouse or trackpad, which triggered the appearance of a randomly oriented white arrow within the central annulus. Observers were instructed to align the direction of the arrow with the previously presented motion direction of the cued stimulus. In the online experiment, responses were confirmed by pressing the spacebar, while in the laboratory experiment, they were confirmed by pressing the right mouse button. In all experiments, trials from different experimental conditions were randomly interleaved.

#### Experiment 1: External reward

In Experiment 1, we investigated how external rewards influence motion reproduction precision. To this end, observers received 15 points for reporting a motion direction within 50° of the target direction when the target was of one colour (e.g., green), and 5 points when it was of the other colour (e.g., blue). Responses that were more than 50 degrees from the target direction did not receive any points. The colour associated with high versus low reward was chosen randomly for each observer at the beginning of the experiment. Both stimuli were presented with the same coherence (85%) and no error feedback was provided. Accumulated points were converted to a bonus payment at the end of the experiment, and observers were informed of this at the beginning of the experiment. Overall, they could collect a maximum of one thousand points, which was equivalent to a bonus payment of £1.50. Observers completed twenty practice trials and one hundred experimental trials. The task took approximately 20 minutes to complete. The trials were divided into two equal blocks with a break of at least one minute in between, and the complete testing session lasted approximately 15 min.

#### Experiment 2: Perceived accuracy

Experiments 2a, 2b, and 2c were designed to investigate the role of feedback on the precision of motion reproduction. In all experiments, at the end of each trial, following the response, we presented feedback showing the reported and target motion directions. Unbeknownst to participants, we manipulated the feedback by artificially minifying or magnifying errors for one stimulus colour. All three experiments were identical except for the following differences. In Experiments 2a and 2c, two stimuli were presented simultaneously for 750 ms at two distinct locations, whereas in Experiment 2b, the stimuli were presented sequentially at the same two locations, each for 750 ms. In Experiment 2b, the order of presentation and the colour cues were balanced across conditions. In Experiments 2a and 2b, feedback for one item was minified and feedback for the other item was magnified. In Experiment 2c, observers completed two tasks on separate days: in one task, feedback for one item was minified while feedback for the other item was veridical; in the other task, feedback for one item was magnified while feedback for the other item was veridical. For Experiment 2c, the order of tasks was randomized, as were the colour pairs associated with the manipulations: one task used a green-blue pair, and the other used an orange-magenta pair.

Feedback magnification was done by shifting the presented target motion direction (*θ*^∗^) away from the reported direction 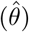 and thereby inflating the presented response error for the designated “difficult” item. This was done according to the following equation:

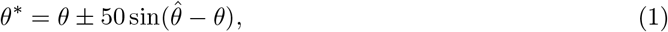

where *θ* is the true motion direction, and all angles are expressed in degrees. Similarly, we systematically minimized the error in the feedback for the other colour, designated as the “easy” item. The magnification and minimization of errors were randomly assigned to one of the two colours (i.e., green or blue, and orange or magenta) for each observer at the beginning of the experiment. The RDK stimuli were presented with 85% coherence. At the beginning of the experiment, during the instructions, we informed observers that individuals might vary in their ability to perceive the motion of stimuli of different colours. This was intended to make any perceived differences in difficulty appear plausible. At the end of the experiment, observers were debriefed and the true purpose of the study was revealed. In all three experiments, observers completed twelve practice trials and one hundred experimental trials. The trials were divided into two equal blocks with a break of at least one minute in between, and the complete testing session lasted approximately 15 min. At the very end of the experiment, observers were asked whether the feedback they received was consistent with their performance and, if not, to elaborate. No participants provided responses indicating that they suspected the feedback was manipulated.

#### Experiment 3: Estimation difficulty

In Experiments 3a and 3b, we investigated the role of stimulus discriminability on the fidelity of visual representations. To achieve this, we presented two stimuli with different levels of coherence on 67% and 70% of all trials in Experiments 3a and 3b, respectively. Specifically, the stimulus of one colour was presented with 85% (high) and the stimulus of the other colour with 45% (low) coherence. These variable-coherence trials were randomly interleaved with trials where both stimuli had the same intermediate (65%) coherence. The assignment of low and high coherence to specific colours was randomized for each observer at the beginning of the experiments. No feedback was provided during these experiments.

Experiment 3a was conducted online, while Experiment 3b took place in the laboratory. In Experiment 3a, on all trials stimuli were presented simultaneously. In Experiment 3b, the same observers performed the task with both simultaneous and sequential presentations, with the order of these conditions counterbalanced across participants. To prevent transfer effects between conditions, we used different colour combinations: in one condition, stimuli were presented in green and blue, while in the other, they were presented in orange and magenta. The colour combinations were randomly assigned to each presentation condition.

In Experiment 3a, observers completed twenty practice trials and one hundred fifty experimental trials. The trials were divided into two blocks with a mandatory break of at least one minute in between, resulting in a total testing session duration of around 15 minutes. Experiment 3b (i.e., the laboratory experiment) consisted of four hundred trials, divided into eight equal blocks. In half of the blocks, stimuli were presented simultaneously, while in the other half, they were presented sequentially. Half of the observers completed the simultaneous blocks first, followed by the sequential blocks, and vice versa for the other half. At the beginning of each block sequence (i.e., simultaneous or sequential task), observers performed twenty practice trials to familiarize themselves with the task. In Experiment 3b, observers were required to maintain central fixation throughout the stimulus presentation. If gaze deviated by more than 2°, a warning message appeared on the screen, and the trial was aborted and restarted with newly randomized stimuli. Completing Experiment 3b took approximately 90 minutes.

### Analysis

All stimulus values were analysed and are reported with respect to the circular parameter space of possible motion directions, [− *π, π*) radians. Response error for each trial was measured as the angular difference between the reported and target motion directions. To quantify the dispersion of response errors, we calculated the mean absolute deviation (MAD) across trials for each condition and observer. Higher MAD values indicate greater average reproduction error. We report all analyses with MAD, but note that we found the same pattern of differences across experiments with other measures of response variability (circular standard deviation and variance, and cosine dissimilarity).

To compare differences in performance across conditions, we used Bayesian hypothesis tests, implemented in JASP [99] with the default Jeffreys-Zellner-Siow prior on effect sizes [100]. We report Bayes factors which compare the relative predictive adequacy of two competing hypotheses (e.g., alternative and null) and quantify the change in belief that the data bring about for the hypotheses under consideration [101]. For example, BF_10_ = 10 indicates that the data are ten times more likely to occur under the alternative hypothesis (i.e., there is a difference) than under the null hypothesis (i.e., there is no difference). Evidence for the null hypothesis is indicated by BF_10_ < 1, in which case the strength of evidence is indicated by 1*/*BF_10_. Evidence assessed via the Bayes Factor is best understood as a ratio-scaled value ranging from 0 to infinity. For clarity in communication, we also use an interpretative framework for Bayes Factor values, following the classification scheme outlined by Lee and Wagenmakers [102]: BF = 1 as no evidence; 1 < BF < 3 as weak or anecdotal evidence; 3 ≤ BF < 10 as moderate evidence; 10 ≤ BF < 30 as strong evidence; 30 ≤ BF < 100 as very strong evidence; BF ≥ 100 as extreme evidence. It is critical to note that while we utilize these discrete categories, they are arbitrary and should serve only as rough guidelines. Along with the Bayes factor, we report the median of the posterior distribution over the effect size (*δ*) and the accompanying 95% credible interval (95% CI).

### Neural resource model

We analysed observers’ response errors with an established model based on the principles of population coding (Fig. 1F; [22, 56, 103, 104]). In this framework, a visual stimulus (*θ*) is encoded by an idealized population of neurons whose activity is determined by their individual tuning functions. All neurons are assumed to share the same bell-shaped von Mises tuning function,

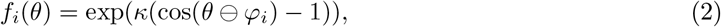

where *κ* determines the tuning concentration, and ⊖ is subtraction on a circle. These tuning functions are translated through the feature space to peak at each neuron’s preferred value (*φ*_*i*_), such that they provide dense uniform coverage of the entire feature space. In a population of *M* neurons encoding *N* visual stimuli, the average response of the *i*th neuron in response to a stimulus value *θ* is:

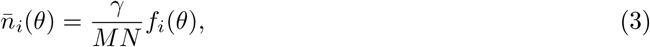

where *γ* is the population’s mean total firing activity (*γ*). This can be simplified by defining 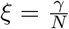 as the mean firing activity encoding a specific stimulus *θ*. If activity associated with multiple stimuli is combined or normalized [105] at a population level *γ*, Equation 3 implements a form of limited resource [22]. The spike count produced by each neuron is drawn from a Poisson distribution,

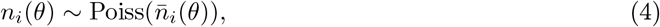

and the decoded motion direction estimate is obtained by maximum likelihood estimation of the population spiking activity, **n**:

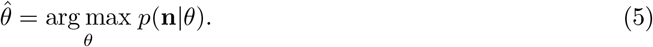

The resulting distribution of decoding errors, for a given total number of spikes *m* = Σ_*i*_*n*_*i*_ ∼ Poiss(*ξ*), is described as a mixture of von Mises (*ϕ*) distributions,

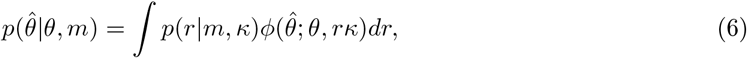

with

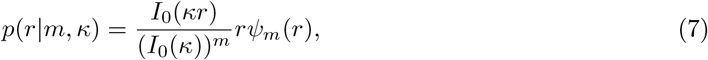

where *rψ*_*m*_(*r*) is the probability density function for resultant length *r* of a uniform random walk of *m* steps. The full distribution of response errors predicted by the model is a mixture of probability distributions 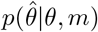, weighted with the probability of obtaining *m* spikes. For a complete derivation of the distribution of response errors, see [22, 75].

While the neural resource model is rooted in principles of neural coding, its predictions for behaviour have been shown to be mathematically equivalent to estimation based on stochastic sampling [56]. According to this interpretation, an estimate is generated by averaging a set of noisy samples of a stimulus; the number of samples varies randomly around a mean determined by the fraction of resources allocated to it. See [56] for further details.

The model has two free parameters, the population’s mean total firing activity (*γ*), and the concentration of the tuning function (*κ*). In scenarios when multiple objects (*N*) need to be represented, the total resource *γ* is typically divided equally among objects (i.e., *ξ* = *γ/N*). Here we extend this basic approach by incorporating an allocation parameter, or gain factor *α*, which controls the neural activity allocated to one object (see also [22]). Without loss of generality, we fixed the gain factor for one object at 1, while treating the gain factor for object *j* (see below for details of each experiment) as a free parameter when fitting the model to the data. The neural activity allocated to object *j* can be expressed as *ξ*_*j*_ = *p*_*α*_*γ*, where *p*_*α*_ = *α/*(1 + *α*) represents the proportion of total neural activity. The remaining activity (proportion 1 − *p*_*α*_) is allocated to the other object.

In Experiment 1, which involved the manipulation of external reward, the allocation weight for the high-reward item was fixed at 1, while the allocation weight for the low-reward item was freely estimated. In Experiment 2, in which we manipulated perceived accuracy, the allocation weight for the error-magnified item was fixed at 1, and the allocation weight for the error-minified item was freely estimated. In Experiment 3, which involved estimation difficulty manipulation, we simultaneously fitted responses on variable- and equal-coherence trials. Building on our previous work [106], we assumed that the strength of the motion signal is controlled by the coherence level of RDK stimuli, such that the value encoded into the neural population is given by

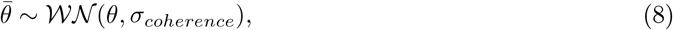

where 𝒲𝒩 is a wrapped normal with mean *θ* and variance *σ*_*coherence*_ accounting for additive Gaussian noise. For simplicity, we considered 85% coherence (high coherence) as perceptually noiseless and further assumed that *σ*_45%_ *> σ*_65%_, where 45% was the low-coherence level, and 65% was the intermediate-coherence level used in the equal-coherence trials. Additionally, the allocation weight for the low-coherence colour (45% and half of 65% stimuli) was fixed at 1, while the allocation weight for the high-coherence colour (85% and half of 65% stimuli) was freely estimated across variable- and equal-coherence trials. In other words, on equal-coherence trials, differences in response precision were explained solely by the allocation weight. In contrast, on variable-coherence trials, the allocation weight and perceptual noise jointly accounted for variations in response precision.

#### Optimal resource allocation

To identify the optimal levels of resource allocation, we conducted a simulation study. For each observer, we simulated model predictions using the best-fitting parameters of the Neural resource model, specifically the population’s mean total firing activity (*γ*), and the concentration of the tuning function (*κ*), along with a grid of potential allocation weights.

For Experiment 1, we analytically determined the number of points based on the model-predicted response distributions under different allocation weights. The allocation weights were tested across a grid ranging from 0.001 to 2 in increments of 0.01, resulting in 200 distinct values. The optimal allocation weight was identified as the value that maximized the total reward across both high- and low-reward items.

For Experiment 2, we numerically simulated the variance (i.e., squared circular SD) of feedback errors using the same grid of allocation weights employed in Experiment 1. This analysis was based on 10^7^ simulated trials drawn from the error distribution predicted by the model. The optimal allocation weight was determined as the value that minimized the total variance of feedback errors across both error-minified and error-magnified items.

For Experiment 3, we analytically modelled the response variance on the variable coherence trials (i.e., for the high- and low-coherence items) across a range of allocation weights. We employed a grid of 200 values, spanning from 0.01 to 6 in increments of 0.03. The optimal allocation weight was identified as the value that minimized the total response variance for both high- and low-coherence items (i.e., 85% and 45% coherence). In Experiment 3a, the simulation yielded values around *α*_optimal_ = 1 for all but one outlier observer, for whom the estimate reached the endpoint of the examined grid (*α*_optimal_ = 6). This occurred due to the model estimating high levels of perceptual noise for medium- and low-coherence stimuli, suggesting that minimizing overall error would be achieved by allocating all resources to the high-coherence object. We exclude this data point in Figure 4D, and the comparison of observed and optimal allocations is based on the remaining observers. Including this observer’s data and performing a non-parametric test did not change our conclusions.

Optimal allocation in this case depends on two additional features of the experiment: first, that the relative frequency of conditions is equal (i.e., an equal number of trials across conditions), and second, that observers cannot predict which item will be probed for reproduction (i.e., trials were randomly interleaved and probed across conditions).

### Reinforcement learning account of resource allocation

We developed a quantitative model to describe how the history of accumulated rewards from multiple objects influences subsequent resource allocation towards those objects. The proposed model extends the Neural resource model by incorporating a simple reinforcement learning (RL) rule, which directs behaviour towards more rewarding stimuli. Importantly, our model applies the same RL rule to both external and intrinsic rewards. In the standard RL framework, analysis typically focuses on external rewards, such as points or money, which are provided by the environment as a direct response to the agent’s actions. Our model broadens this scope to include intrinsic rewards – those that are inherently pleasurable and drive behaviour – such as the sense of being accurate in a task.

Drawing on the conducted experiments and the motion direction reproduction task, the general overview of this account is as follows: on a particular trial, the received points or money (external reward), perceived accuracy due to feedback (intrinsic reward), and an individual’s internal confidence-based estimate of precision (intrinsic reward) collectively update the value (*ν*) of a particular object associated with these rewards. This computed value influences the allocation of cognitive resources to that object in subsequent encounters, thereby modulating the precision with which the object is represented.

Formally, in the simplest scenario involving only two objects, rather than defining the accumulated reward for each object separately, we can define the relative accumulated reward on trial *t* as:

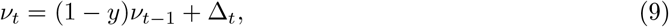

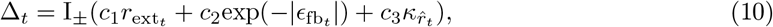

where *y* is a leak component, 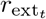 is the number of received points, 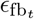 is feedback error, 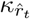 is an estimate of internal confidence, and {*c*_1_, *c*_2_, *c*_3_} are respective weights accounting for different scales of rewards and the types of rewards prioritised by observers. The variable I_*±*_ takes the value of +1 or -1 depending on the object identity, i.e., reproduction of the green or blue item, with the assignment of conditions being arbitrary. Positive values of *ν* indicate a higher relative value for one item (I = +1), while negative values of *ν* indicate a higher relative value for the other item (I = − 1). To account for the fact that *ν* can range from −∞ to +∞, we transform it into gain parameter *α* using the following equation:

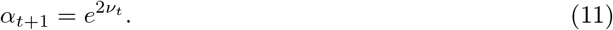

This allows us to compute the proportion of spiking activity that will be allocated on a subsequent trial (*t*+1) to the item identified as sign(*I*) = +1, given by *α/*(1 + *α*), with the remaining spiking activity allocated to the other item. When *ν* = 0, such as at the beginning of the task, both items are perceived as having equal value, resulting in an equal distribution of neural resources between them.

The leak component (*y*) functions as a temporal filter, modulating the influence of past rewards on resource allocation. When *y* = 1, the system entirely ignores accumulated past values, making the value of an object – and thus resource allocation in the next trial – rely exclusively on the reward from the most recent trial. Conversely, when *y* = 0, the accumulated value is fully retained and integrated with the most recent reward. The necessity of the leak component becomes particularly evident in scenarios where rewards are discontinued: a non-zero leak will gradually equalize the relative value and resource allocation across objects, returning them to a state of equilibrium.

The first reward component of Equation 9, *r*_ext_, reflects the experimental manipulation of Experiment 1. In this experiment, observers received 15 points for responses with an error of less than 50° for high-reward objects and 5 points for low-reward objects. Responses with an error greater than 50° received no points. When applying this model to the data, we used values of *r*_ext_ = {0.15, 0.05, 0 } to represent the rewards for high-reward, low-reward, and no-reward trials, respectively. Note that the utility of rewards, and especially money, is often considered a nonlinear function rather than linear, as assumed here [107, 108]. We employed a linear utility function for simplicity and mathematical convenience. Incorporating a parametric non-linear utility function (e.g., a power law with exponent *β*) would introduce an additional free parameter without changing the qualitative pattern of predicted results or the main conclusion: that resource allocation is biased towards items providing higher rewards. Future studies could explore conditions in which non-linear reward functions, reflecting the concave utility of money, might prove critical for explaining behaviour in similar psychophysical tasks.

The feedback component of the model (*ϵ*_fb_) addresses the experimental manipulation of Experiment 2. In this experiment, we systematically manipulated feedback error by reducing it for one stimulus and increasing it for another. We hypothesized that feedback serves as an intrinsic reward, with stimuli receiving minified feedback errors being perceived as more rewarding than those with magnified feedback errors. In modelling this relationship, feedback error was assumed to be exponentially related to the object’s value *ν*, with smaller feedback errors corresponding to higher rewards, leading to a greater increase in the object’s value. This exponential relationship reflects diminishing sensitivity to large feedback errors, such that a wide range of larger errors yields relatively minimal and similar rewards, whereas a narrow range of smaller errors results in significantly higher but more variable rewards.

The final component of our model is the estimate of internal confidence 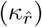. While internal confidence can be assessed through self-reported or metacognitive measures, our approach leverages the inherent mechanism of the Neural resource model to quantify uncertainty in the decoded (i.e., reported) value. Our approach relies on the principle that the width of the likelihood function reflects the uncertainty of the estimate. The likelihood function evaluates how well various stimulus values align with the observed neural activity: a broad likelihood function is compatible with many different feature values, suggesting lower precision in the maximum likelhood estimate (the peak of the likelihood function), whereas a narrow likelihood function implies a more precise estimate. Due to the probabilistic generation of spikes across retrievals (Eq.4), the likelihood has the form of a von Mises with concentration 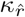 proportional to the resultant vector length of the preferred values associated with each of the emitted spikes (*m*), with higher spike counts producing a narrower likelihood function on average [56]. This formulation has previously been shown to quantitatively reproduce findings from studies in which participants were asked to rate their subjective confidence in each estimate [75, 78]. Consequently, the precision of the likelihood function emerges as a natural candidate for a computational estimate of the observer’s internal confidence.

To measure internal confidence associated with each response, we determined the most probable resultant vector length given the individual response errors and the probabilistic distribution of spike values, which was fully characterized by *ξ* and *κ*. Specifically, for each trial, we used Bayes rule to find the posterior probability of resultant vector length *r* given the error on that trial *ϵ*, marginalizing over total spike count *m*,

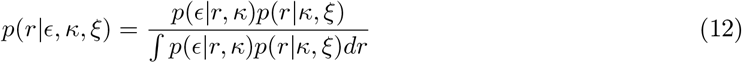

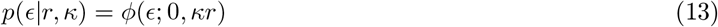

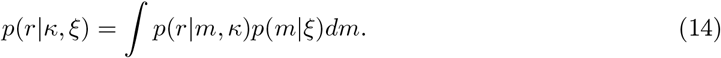

where *p*(*r*|*m, κ*) is given by Eq. 7 and *p*(*m*|*ξ*) is the Poisson p.m.f. with mean *ξ*. Applying MAP estimation to this posterior distribution returns the most probable estimate of resultant length for a given response error,

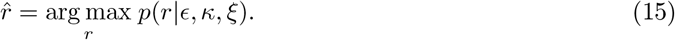

We then use 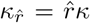 as a measure of internal confidence on the given trial. For experiments with variable estimation difficulty (Experiment 3) the pdf *p*(*ϵ*|*r, κ*) was obtained by convolving the von Mises pdf (the RHS of Eq. 13) with a wrapped normal distribution with freely fitted standard deviation. Finally, 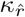 was then calculated as the concentration parameter of a von Mises distribution whose resultant length matches that of 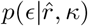.

### Model fitting

To model the observed allocation within the Neural resource model [22, 56], which has two free parameters – the mean population activity (*γ*) and the precision of the tuning functions (*κ*) – we introduced an additional parameter, the gain modulation *α* [22], resulting in a total of three free parameters in Experiments 1 and 2. In Experiment 3, which involved an estimation difficulty manipulation, the Neural Resource model was extended by two additional parameters (*σ*_45%_ and *σ*_65%_) to capture the effects of variable sensory noise introduced by different coherence levels. This brought the total number of free parameters in the Neural resource model for Experiment 3 to five.

The Reinforcement learning account retained all parameters of the Neural resource model except the gain modulation parameter *α*, while introducing four new parameters, namely the leak parameter (*y*) and reward weight parameters (*c*_1_, *c*_2_, *c*_3_). In all three experiments, we modelled the leak parameter (*y*) and the effect of internal confidence (*c*_3_) on resource allocation (see Eq. 9); additionally, we modelled the effect of external reward (*c*_1_) only in Experiment 1 while setting it to zero in all other experiments, and feedback error (*c*_2_) only in Experiment 2 while also setting it to zero in all other experiments. This resulted in the estimation of five free parameters in Experiments 1 and 2, and six in Experiment 3. When fitting the model to the data, the leak parameter was constrained between 0 and 1, and all three weight parameters were limited to a range of -1 to 1.

For all models, we obtained a separate maximum likelihood fit for each individual observer. These fits were derived using the Nelder-Mead simplex method (via the *fminsearch* function in MATLAB). A MATLAB toolbox implementing the Neural resource model is available for download from https://bayslab.com/toolbox.

## Acknowledgment

We thank David Aagten-Murphy and Robert Taylor, who worked on earlier iterations of this project. We thank Neha Abraham, Pepita Alex, Amida Anand, Paul McMeekin, Adam Sabo, Tom Wenban-Smith, Adam Zhu, and Adam Triabhall for assisting with data collection.

We used resources provided by the Cambridge Service for Data Driven Discovery (CSD3) operated by the University of Cambridge Research Computing Service.

This work was funded by the Wellcome Trust (grant 106926 to P.M.B). The funders had no role in study design, data collection and analysis, decision to publish or preparation of the manuscript.

## Author contributions

**I.T**. contributed to conceptualization, methodology, software, data collection, investigation, formal analysis, modelling, visualizations, and writing – original draft and revisions. **R.R.R** contributed to methodology, software, and data collection. **P.M.B**. contributed to conceptualization, funding acquisition, supervision, methodology, formal analysis, modelling, visualizations, and writing – editing and revisions.

## Data and code availability statement

Data and analysis code will be made publicly available upon publication of this manuscript, with a DOI-assigned version deposited in a publicly accessible repository.

## Supplementary Information

### Psychophysical data

#### Experiment 2b: Sequential presentation

**Figure S1:**
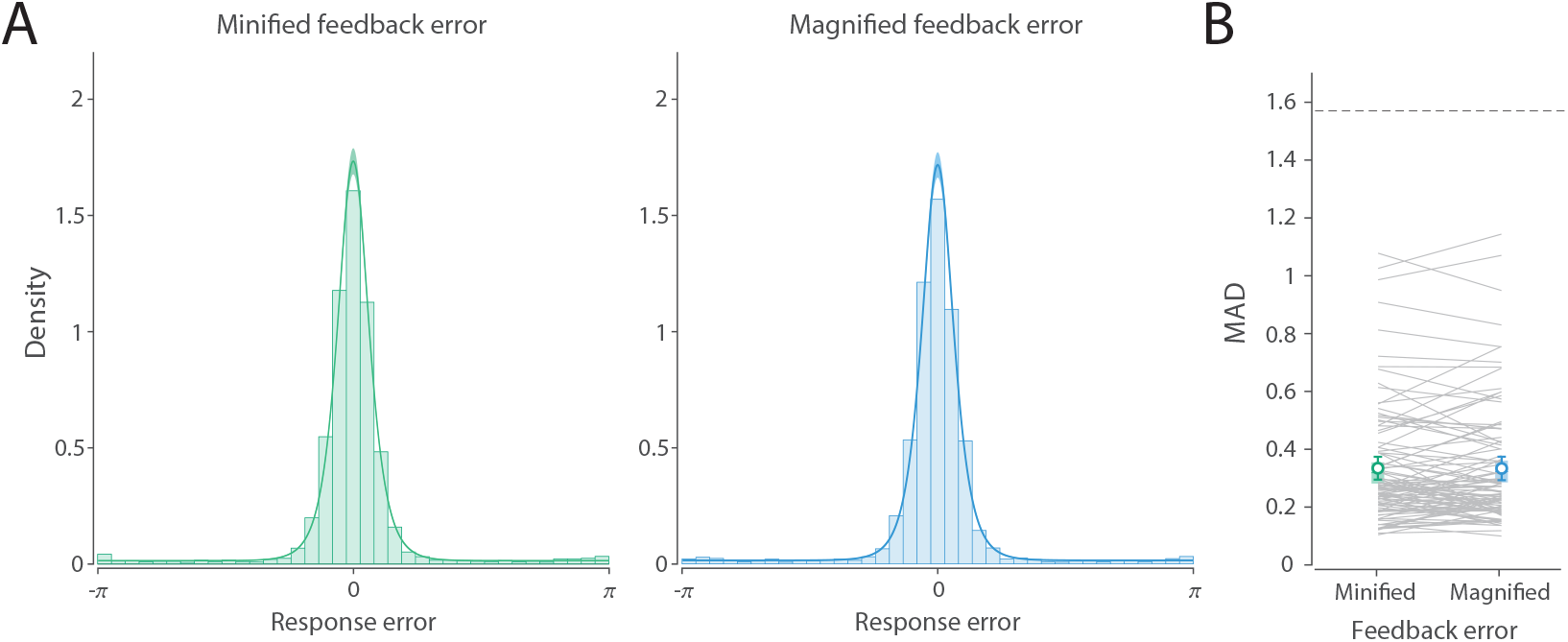
Perceived accuracy manipulation in Experiment 2b (sequential presentation). A) Distribution of response errors and corresponding fits of the Neural resource model. Histograms represent the data, while coloured curves and shaded areas depict model predictions (M ± SE). B) Corresponding MAD from experimental data (circles with error bars) and from the Neural resource model (lines with error patches). Error bars and patches indicate the mean and 95% CI. Dashed line indicates chance level performance.

#### Experiment 3b: Sequential presentation

**Figure S2:**
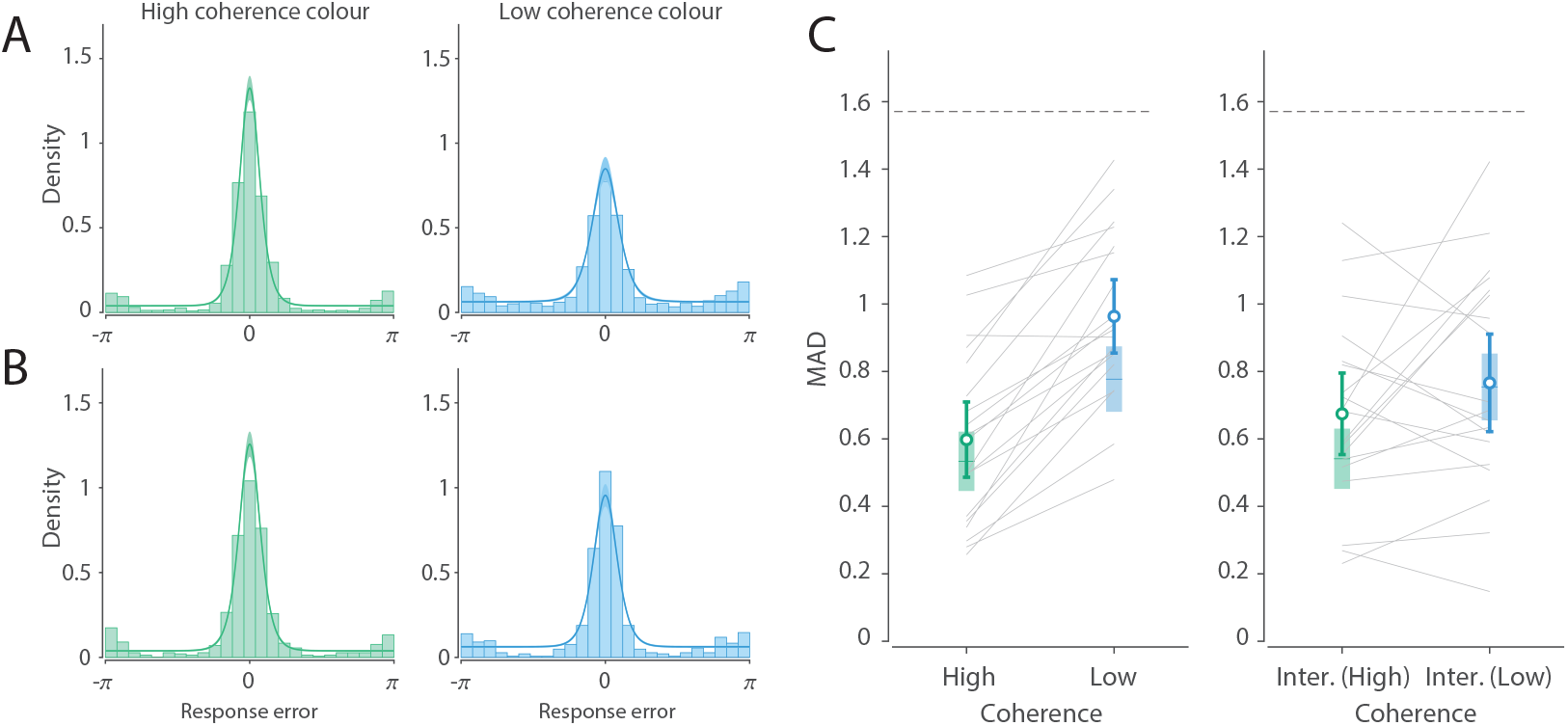
Estimation difficulty manipulation in Experiment 3b (sequential presentation). A & B) Histograms represent distributions of response errors, while coloured curves and shaded areas depict model predictions. Panel A depicts variable coherence trials, and panel B depicts equal coherence trials. C) Corresponding MAD from experimental data (circles with error bars) and from the Neural resource model (lines with error patches). Error bars and patches indicate the mean and 95% CI. Dashed line indicates chance level performance.

### Neural resource model

#### Parameter recovery

To test the identifiability of model parameters, we performed parameter recovery. To this end, we pooled (static) Neural resource model parameters estimated in Experiments 1 and 2a, resulting in 55 unique parameter sets (Experiments 3a and 3b produced very similar average estimates to these and were therefore not included here). For each set of parameters (i.e., each observer), we simulated 20 datasets of the same size as the individual datasets reported in the paper (i.e., 100 trials in total). This procedure necessarily introduces some randomness in the recovered parameters. We then performed the model fitting procedure as described in the paper. To ensure that we found the global minimum, we initiated the parameter estimation procedure from multiple starting points, as we did for our main analysis. In Fig. S3, the confusion matrices show the distributions of recovered parameters for each set of ground truth parameters. For ease of visualization, we normalized all probabilities column-wise, i.e., within each bin of ground truth parameters. The identity line indicates perfect recovery, which would be achieved with an infinite amount of data. In all cases, the recovered parameters are strongly clustered around the ground truth values.

**Figure S3:**
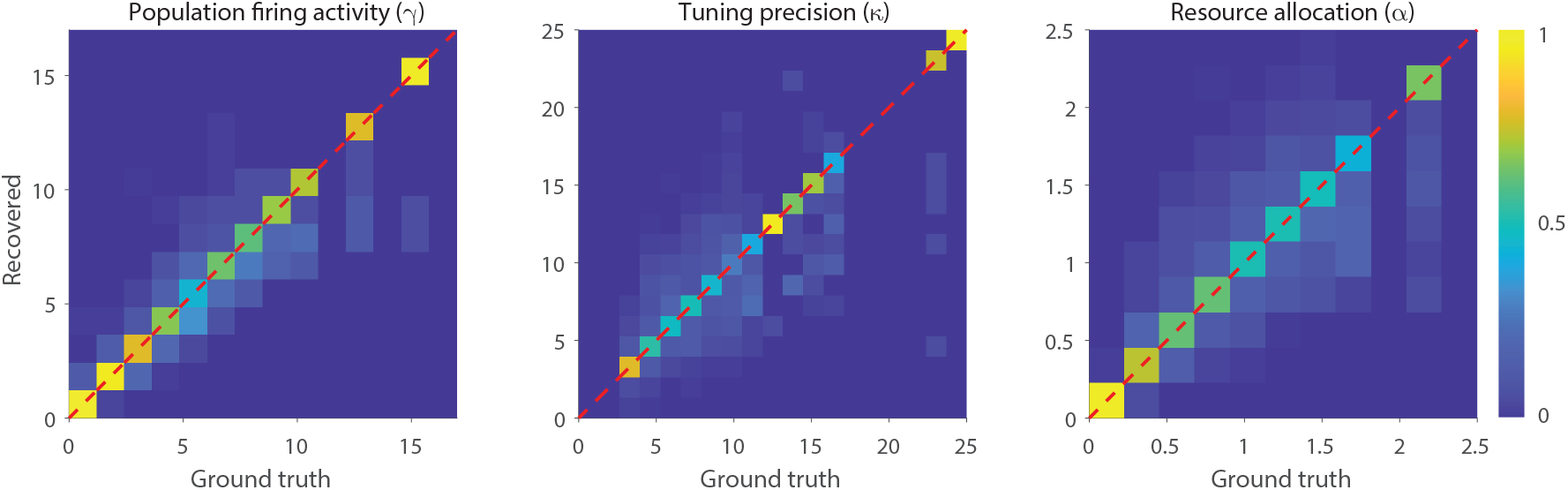
Parameter recovery for population firing activity, tuning precision, and resource allocation parameters. Dashed diagonal (i.e., identity) line indicates perfect recovery.

#### Optimal allocation under different loss functions

To derive the optimal allocation for Experiments 2 and 3, we assumed that optimal behaviour corresponds to minimizing the expected squared error – the feedback error when available (Experiment 2) or else the response error (Experiment 3). To test the generality of our findings, we simulated optimal allocations under alternative loss functions. Specifically, we defined loss as *L*_*p*_ = (1 − *cos*(*error*))^(*p/*2)^, where *p* is the loss exponent (*p >* 0). Larger values of *p* penalize large errors more strongly. The definition of optimality in the main analysis is approximately equivalent to the allocation that minimizes *L*_2_. In Experiment 2a, for all integer loss exponents, the optimal resource allocation was strongly biased towards the error-magnified object (Fig. S4A). In Experiment 3a, the optimal resource allocation was approximately equal in every case (Fig. S4B). In other words, for both linear (*p* = 1) and supralinear (*p >* 1) loss exponents, the predictions regarding optimal allocation were qualitatively similar and differed strongly from the observed allocation, as in the main analysis.

We also considered the hit-or-miss (*L*_0_) loss function, where all non-zero errors incur equal loss: in this unusual case the optimal strategy was to allocate all resources to the “easier” item (i.e., minified feedback or high coherence), equivalent to *α* = ∞. This allocation is also inconsistent with the observed results. While it is sometimes used for scientific computation and model fitting, *L*_0_ is not generally considered a plausible behavioural loss function, because zero error can never be achieved in real scenarios. Previous studies suggest that human errors across a range of tasks are consistent with a supralinear loss function, *p* = 2 or greater [22, 25, 109–111].

#### Parameter estimates from the Neural resource model

Applying the static version of the Neural resource model to experimental data, in addition to the values of the *α* parameter presented in the main text, we observed the following ML estimates for the population’s mean total firing activity and concentration of the tuning function (mean ± SE): Experiment 1: *γ* = 2.88 ± 0.40, *κ* = 10.14 ± 0.91; Experiment 2a: *γ* = 4.81 ± 0.69, *κ* = 6.83 ± 0.55; Experiment 3a: *γ* = 3.39 ± 0.40, *κ* = 10.55 ± 0.65; Experiment 3b: *γ* = 2.11 ± 0.31, *κ* = 12.07 ± 1.30.

**Figure S4:**
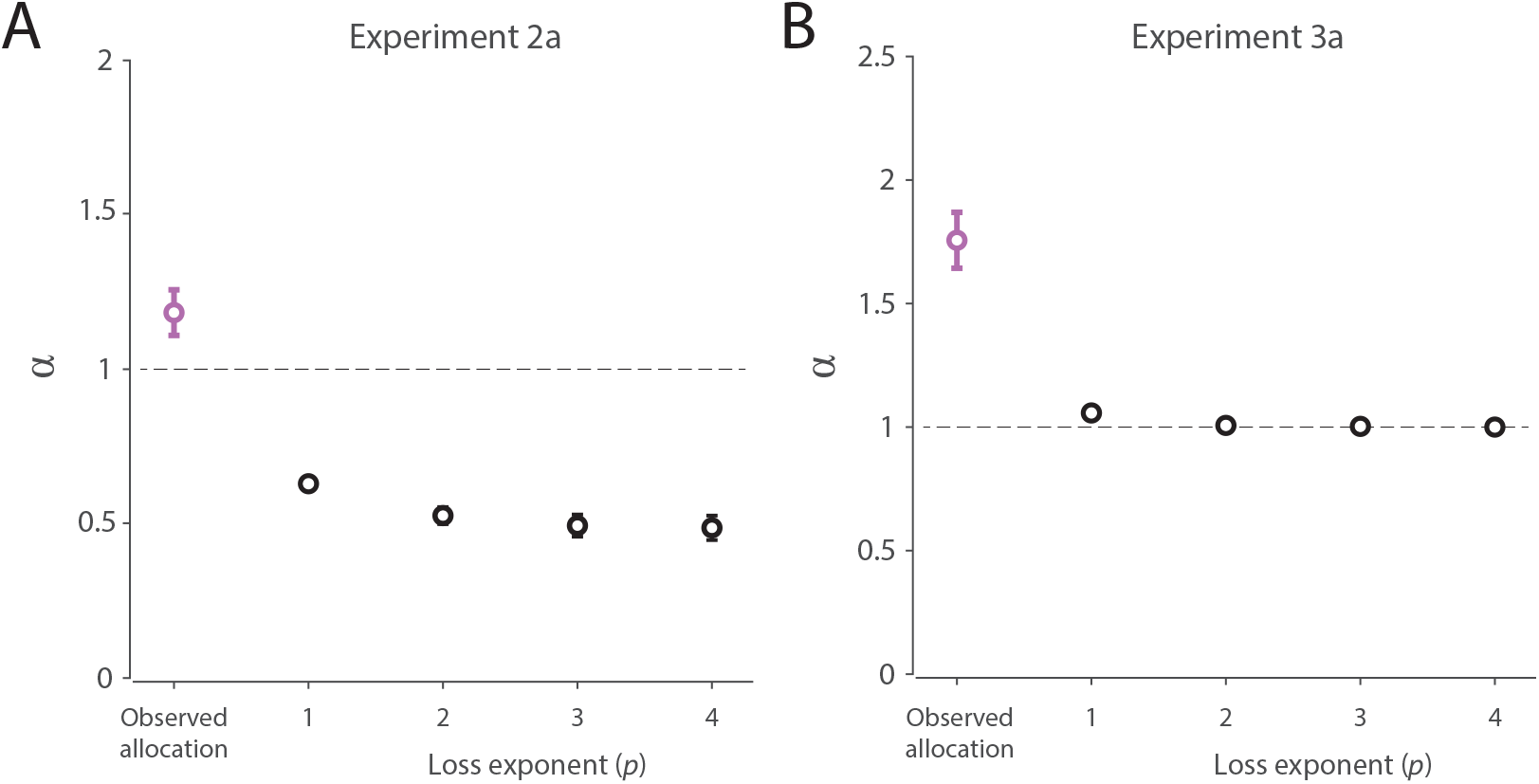
Optimal allocation under loss functions identified by different loss exponents. A) In Experiment 2a, linear (*p* = 1) and supralinear (*p >* 1) loss functions predict an optimal allocation of resources that strongly favours the error-magnified item (*α <* 1). B) In Experiment 3a, all tested exponents predicted approximately equal allocation to high- and low-coherence items. In both panels, exponent *p* = 2 is approximately equivalent to the optimal allocation reported in the main text.

Small variations between experiments are due to individual differences between observers, as well as differences in motion coherence between studies, and in online versus lab settings. The observed values are generally very consistent with estimates reported in similar studies [22, 104, 106]. One particular difference worth noting is that estimates of gain parameter are slightly lower, and estimates of concentration parameter are slightly higher for motion stimuli compared to other low-level visual stimuli, such as orientation. This is also consistent with previous observations (e.g., compare Table A1 and Table A3 in McMaster et al. [112].

### Reinforcement learning account

#### External reward

The average trajectory of resource allocation shown in Figure 6A is based on ML parameter estimates (mean ± SE): mean activity *γ* = 2.88 ± 0.39; tuning precision *κ* = 10.29 ± 0.98; leak *y* = 0.28 ± 0.06; reward weight *c*_1_ = 0.31± 0.14; internal confidence weight *c*_3_ = 0.012 ± 0.01. Calculating the correlation between parameter estimates from the Reinforcement learning account and the neural model with freely estimated resource allocation, we found highly consistent estimates of the population’s mean spiking activity (*r* = 0.997, 95% CI = [0.992, 0.999], BF_10_ = 1.46 *×* 10^28^) and tuning precision (*r* = 0.971, 95% CI = [0.929, 0.986], BF_10_ = 3.07 *×* 10^15^) (Fig. S6A).

The full model, incorporating both external rewards and confidence, outperformed a version that included only external rewards (ΔAIC = 62.9), as well as a version that included only internal confidence (ΔAIC = 45.2). Finally, to assess whether observers prioritised external rewards or internal confidence signals in resource allocation, we calculated each observer’s mean contribution of external rewards and internal confidence to the relative value of the two objects. Results indicated moderate evidence for no difference between their contributions (BF_10_ = 0.22; *δ* = 0.076, 95% CI = [-0.263, 0.418]). However, this finding should be interpreted with caution, as the effect of internal confidence may partly reflect observers’ resource allocation favouring the high-reward item (reflecting the influence of external reward), which subsequently enhances confidence for that item.

#### Intrinsic reward: Perceived accuracy

The mean trajectory of resource allocation across trials shown in Figure 6D is based on ML parameter estimates (mean ± SE): mean activity *γ* = 5.01 ± 0.78; tuning precision *κ* = 6.64 ± 0.51; leak *y* = 0.538 ± 0.076; feedback weight *c*_2_ = 0.238 ± 0.108; internal confidence weight *c*_3_ = 0.007 ± 0.044. Comparing the estimates derived from the Neural resource model and the Reinforcement learning account (Fig. S6B), we again found that the RL account’s estimates closely match the population’s mean spiking activity (*r* = 0.987, 95% CI = [0.964, 0.994], BF_10_ = 1.26 *×* 10^16^) and tuning precision (*r* = 0.956, 95% CI = [0.884, 0.980], BF_10_ = 4.7 *×* 10^10^).

As in Experiment 1, the full model incorporating both feedback and internal confidence as sources of reward outperformed the model using only feedback (ΔAIC = 12.1) as well as the confidence-only model (ΔAIC = 7.8). Finally, we found moderate evidence for no difference between feedback and internal confidence signals in their contribution to the relative value of objects (BF_10_ = 0.33; *δ* = 0.174, 95% CI = [-0.196, 0.553]).

#### Intrinsic reward: Estimation difficulty

Fitting the Reinforcement learning account to psychophysical data from Experiment 3a, we obtained the following ML parameters (mean ± SE): mean activity *γ* = 3.33 ± 0.39; tuning precision *κ* = 11.53 ± 0.61; leak *y* = 0.398 ± 0.086; confidence weight *c*_3_ = 0.062 ± 0.045; intermediate perceptual noise *σ*_65%_ = 0.143 ± 0.021; high perceptual noise *σ*_45%_ = 0.338 ± 0.086. In Experiment 3b we observed very similar estimates: mean activity *γ* = 2.13 ± 0.31; tuning precision *κ* = 13.03 ± 1.70; leak *y* = 0.285 ± 0.083; confidence weight *c*_3_ = 0.007 ± 0.004; intermediate perceptual noise *σ*_65%_ = 0.090 ± 0.021; high perceptual noise *σ*_45%_ = 0.249 ± 0.056. Again, we visualised the obtained individual trajectories in example participants (Fig. S5C & D).

In both experiments, estimates obtained with the Neural resource model and the Reinforcement learning account strongly covaried (Fig. S6C & D). Specifically, we found highly consistent estimates of the population’s mean spiking activity (Exp 3a: *r* = 0.995, 95% CI = [0.986, 0.998], BF_10_ = 6.15 × 10^17^; Exp 3b: *r* = 0.999, 95% CI = [0.996, 1.000],BF_10_ = 1.66 × 10^19^), tuning precision (Exp 3a: *r* = 0.912, 95% CI = [0.761, 0.961], BF_10_ = 2.84 × 10^6^; Exp 3b: *r* = 0.970, 95% CI = [0.899, 0.989], BF_10_ = 5.66 × 10^8^), intermediate perceptual noise (Exp 3a: *r* = 0.922, 95% CI = [0.785, 0.966], BF_10_ = 8.00 × 10^6^; Exp 3b: *r* = 0.985, 95% CI = [0.947, 0.994], BF_10_ = 8.17 × 10^10^), and high perceptual noise (Exp 3a: *r* = 0.659, 95% CI = [0.293, 0.830], BF_10_ = 48.1; Exp 3b: *r* = 0.896, 95% CI = [0.696, 0.957], BF_10_ = 6.80 *×* 10^5^).

**Figure S5:**
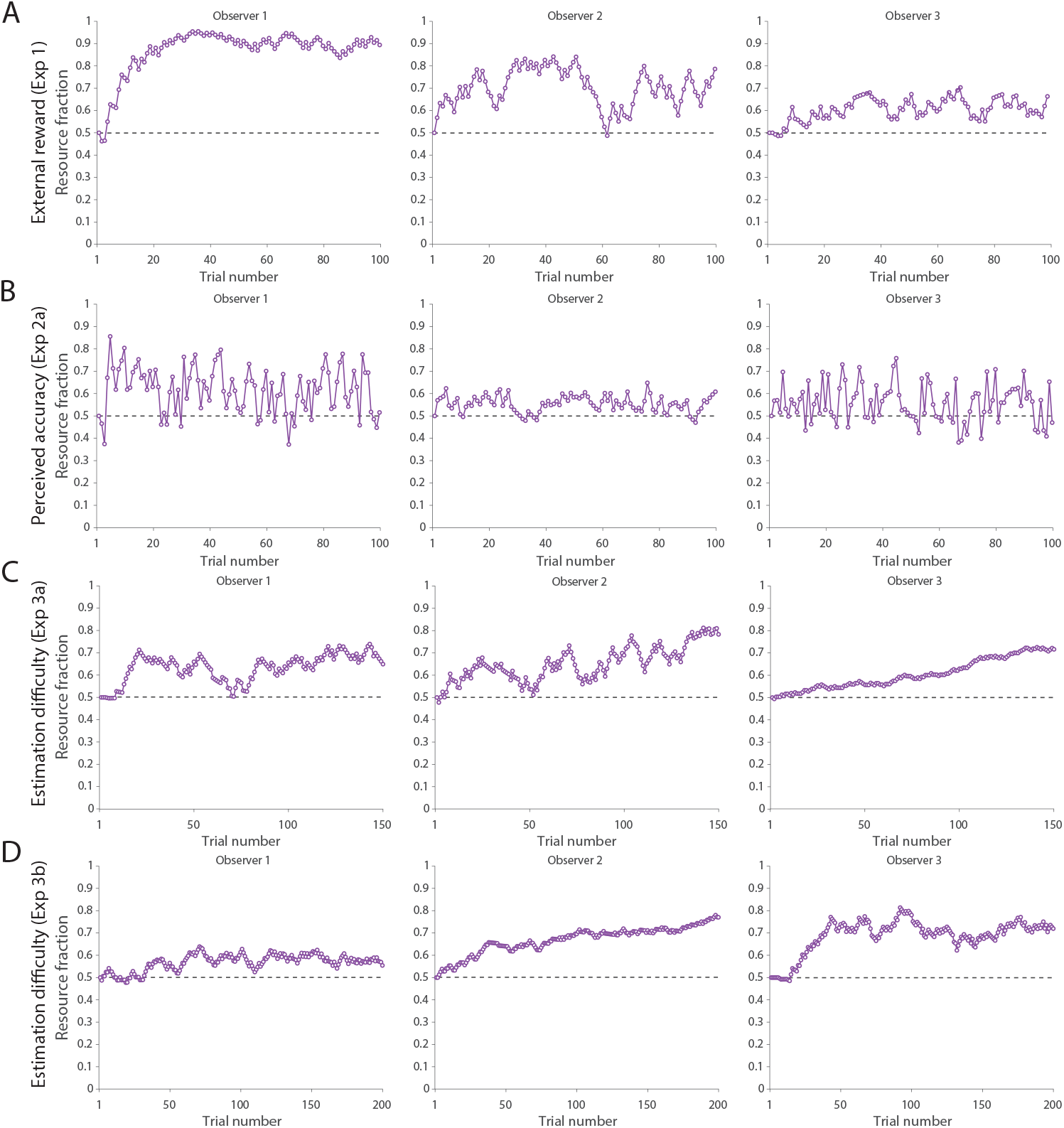
A) Trial-by-trial resource allocation estimated by the RL account in the external reward experiment (Experiment 1) for three illustrative participants. Circles represent the fraction of resources allocated to the preferred item on each trial. B) Perceived accuracy experiment (Experiment 2). C) Estimation difficulty experiment (Experiment 3a). D) Estimation difficulty experiment (Experiment 3b).

**Figure S6:**
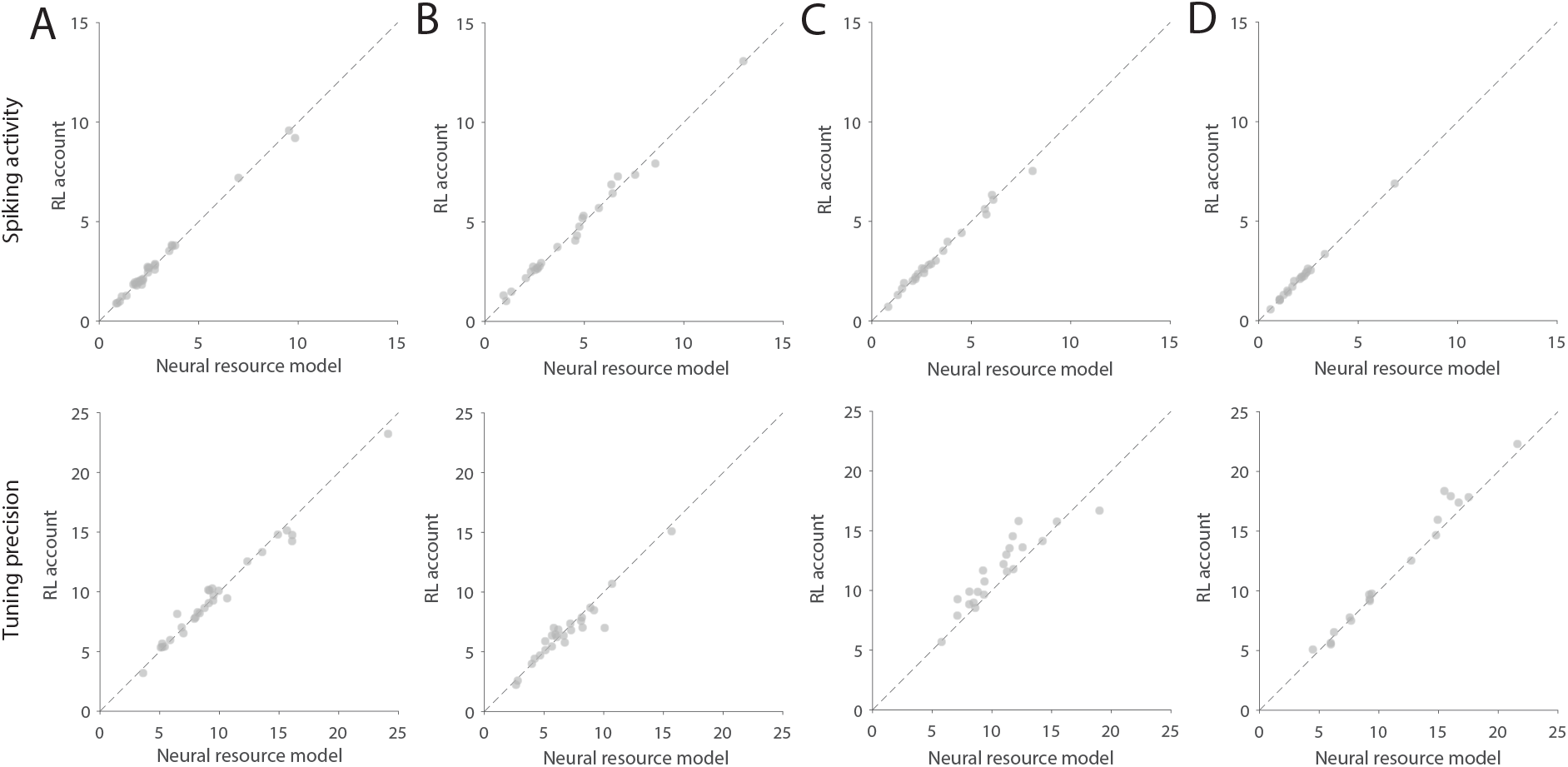
Correlation between mean activity (top row) and tuning precision (bottom row) parameters estimated in the Neural resource model and the RL account of resource allocation. A) External reward experiment (Experiment 1). B) Perceived accuracy experiment (Experiment 2). C) Estimation difficulty experiment (Experiment 3a). D) Estimation difficulty experiment (Experiment 3b).

